# Using virtual patient cohorts to uncover immune response differences in cancer and immunosuppressed COVID-19 patients

**DOI:** 10.1101/2024.08.01.605860

**Authors:** Sonia Gazeau, Xiaoyan Deng, Elsa Brunet-Ratnasingham, Daniel E. Kaufmann, Catherine Larochelle, Penelope A. Morel, Jane M. Heffernan, Courtney L. Davis, Amber M. Smith, Adrianne L. Jenner, Morgan Craig

## Abstract

The COVID-19 pandemic caused by the severe acute respiratory syndrome coronavirus-2 (SARS-CoV-2) resulted in millions of deaths globally. Adults with immunosuppression (e.g., solid organ transplant recipients) and those undergoing active cancer treatments experience worse infections and more severe COVID-19. It is difficult to conduct clinical studies in these populations, resulting in a restricted amount of data that can be used to relate mechanisms of immune dysfunction to COVID-19 outcomes in these vulnerable groups. To study immune dynamics after infection with SARS-CoV-2 and to investigate drivers of COVID-19 severity in individuals with cancer and immunosuppression, we adapted our mathematical model of the immune response during COVID-19 and generated virtual patient cohorts of cancer and immunosuppressed patients. The cohorts of plausible patients recapitulated available longitudinal clinical data collected from patients in Montréal, Canada area hospitals. Our model predicted that both cancer and immunosuppressed virtual patients with severe COVID-19 had decreased CD8+ T cells, elevated interleukin-6 concentrations, and delayed type I interferon peaks compared to those with mild COVID-19 outcomes. Additionally, our results suggest that cancer patients experience higher viral loads (however, with no direct relation with severity), likely because of decreased initial neutrophil counts (i.e., neutropenia), a frequent toxic side effect of anti-cancer therapy. Furthermore, severe cancer and immunosuppressed virtual patients suffered a high degree of tissue damage associated with elevated neutrophils. Lastly, parameter values associated with monocyte recruitment by infected cells were found to be elevated in severe cancer and immunosuppressed patients with respect to the COVID-19 reference group. Together, our study highlights that dysfunction in type I interferon and CD8+ T cells are key drivers of immune dysregulation in COVID-19, particularly in cancer patients and immunosuppressed individuals.

## INTRODUCTION

The COVID-19 pandemic caused by the severe acute respiratory syndrome coronavirus-2 (SARS-CoV-2) caused more than 7 million deaths globally as of July 2024^1^. COVID-19 results in heterogeneous immune responses and outcomes, where some individuals experience no or very few symptoms while others become hyperinflamed and may need supportive oxygen or succumb to the infection. The risk of severe complications to SARS-CoV-2 infection is increased for individuals with weakened or suppressed immune responses^2^. Thus, it is critically important to study immuno-infection dynamics, especially in vulnerable groups (e.g., immunocompromised individuals^3–5^ such as patients receiving immunosuppressants after organ transplantation, cancer patients^6–8^, older adults^9,10^) whose immune systems may not adequately protect against the virus and who may have imperfect vaccine-induced immune responses^11^.

Cancer patients tend to have weaker responses to viral infections^7^, mostly due to impaired responses of type I interferon (IFN) that are typical of cancers^12^ and diverse immune cell dysfunctions that are frequent adverse effects of oncologic treatments. COVID-19 mortality risk in patients with hematological malignancies is around 34%, although a study by Vijenthira et al. found the most relevant factor impacting mortality to be age^13^. COVID-19-positive leukemia patients have an increased fatality rate compared to the patients with other cancer types^14^, likely due to the susceptibility of blood cancer patients to experience lymphocyte depletion. Furthermore, anti-cancer treatments like cytotoxic chemotherapy can result in decreased T and B lymphocytes^15,16^, leaving patients undergoing treatment vulnerable to severe infections. It has also been observed that patients receiving anti-cancer therapy tend to have low platelet and/or decreased neutrophil counts^15^. Although some studies have reported neutropenia as a risk factor in COVID-19-positive hematological malignancy patients^16^, others found no significant connection^17^. However, cancer patients often have hyperactivated IL-6^5,7^, which may be another factor affecting COVID-19 severity, as multiple studies have shown that elevated IL-6 concentrations are associated with poor COVID-19 outcomes^18–20^.

Immunosuppressed patients, such as solid organ transplant recipients, are treated with anti-T or anti-B cell therapies to prevent immunological rejection of transplantable tissue. This results in decreased lymphocytes^3^. A recent study showed that IL-6 concentrations in COVID-19 immunosuppressed patients without autoimmune disease were significantly increased compared to COVID-19 patients without immunosuppression^4^. Together, this lack of lymphocytes and elevated concentrations of inflammatory cytokines (i.e., IL-6) results in a weakened immune response against acute infections including ones caused by respiratory viruses, resulting in severe infections^21^. IL-6 dysregulation may result in hyperinflammation that is characteristic of severe COVID-19, particularly in patients requiring intensive care^22^.

Once it was identified that extreme inflammatory responses could develop from SARS-CoV-2 infections, potential causes and treatment strategies were intensively studied^22–24^. Although the direct causes of hyperinflammation have yet to be established, several hypotheses exist^22^. One links the condition with the viral replication leading to pyroptosis, a highly inflammatory form of apoptosis, which then causes a pro-inflammatory cytokine reaction that affects macrophages and lymphocytes^22^ and causes excessive IL-6 production^25^. Others include uncontrolled adaptive and neutralizing antibody responses, proposing that antibody binding to spike protein causes hyperinflammation^22^. Collecting longitudinal data in humans, particularly vulnerable populations, can be difficult and thus limited in scope. Further, these studies may not be able to uncover kinetic differences and causes for dysregulated immune responses, which are difficult to establish in humans, particularly given that early infection dynamics are generally not captured in clinical data. Mechanistic mathematical modelling helps to overcome these complexities because it allows for the investigation of immune response mechanisms, aids the prediction of clinical outcomes or vaccination efficacy^26,27^, and facilitates uncovering potential drivers of severity with limited data sources^28^.

Here, we focused on investigating and predicting COVID-19 immune dynamics in vulnerable populations, including those undergoing cancer treatments or who are immunosuppressed without autoimmune disease (e.g., solid organ transplant recipients on immunosuppressive agents). For this, we extended our approach described in Jenner et al.^28^, where we generated a cohort of COVID-19-positive virtual patients based on a mechanistic model of the immune response to SARS-CoV-2. The model predicted that patients with severe outcomes are more likely to experience delayed IFN peaks and CD8+ T cell depletion. Because our previous work did not take existing comorbidities into account, in this study we used the same model to generate three virtual patient cohorts: 1) a cohort of COVID-19+ patients with cancer, 2) a cohort of COVID-19+ immunosuppressed patients, and 3) a reference group of COVID-19+ patients without cancer or immunosuppression. The virtual patient cohorts were based on data collected from Montréal, Canada area hospitals^29–31^ and data available in the literature. Our simulations suggested that both severe cancer and immunosuppressed patients have decreased CD8+ T cells, elevated neutrophils and IL-6 concentrations, and delayed IFN peaks. As in our previous work, we found these alterations to be driven by monocyte to macrophage differentiation and monocyte recruitment, consistent with experimental and clinical studies^32–34^, suggesting these are host-intrinsic rather than driven by comorbidities. Overall, our findings suggest suppressed CD8+ T cells, overproduction of IL-6, and delayed IFN peaks are correlated with disease severity in cancer and immunosuppressed patients with COVID-19, similar to previous results in COVID-19 severe virtual patients described in Jenner et al.^28^ However, we determined that the most severe outcomes in cancer and immunosuppressed virtual patients were characterized by more marked increases in elevated neutrophils during infection, higher rates of monocyte to macrophage differentiation by IL-6, and increased monocyte recruitment by infected cells. Thus, our study further highlights that immune dysfunction is heightened in immunocompromised patients, with potential consequences on COVID-19 severity, and identifies biomarkers driving this dysregulation.

## METHODS

### Mathematical model of the immune response to SARS-CoV-2

We used the differential equation-based mathematical model of Jenner et al.^28^ that mimics the immune response to SARS-CoV-2 to understand and predict immune dynamics during COVID-19 (Figure 1A). The model describes the dynamics of immune cells (neutrophils, monocytes, CD8+ T cells, and tissue-resident and inflammatory macrophages) together with cytokine production and binding kinetics, including IFN-α,β, IL-6, granulocyte-macrophage colony-stimulating factor (GM-CSF), and granulocyte colony-stimulating factor (G-CSF).

**Figure 1.**
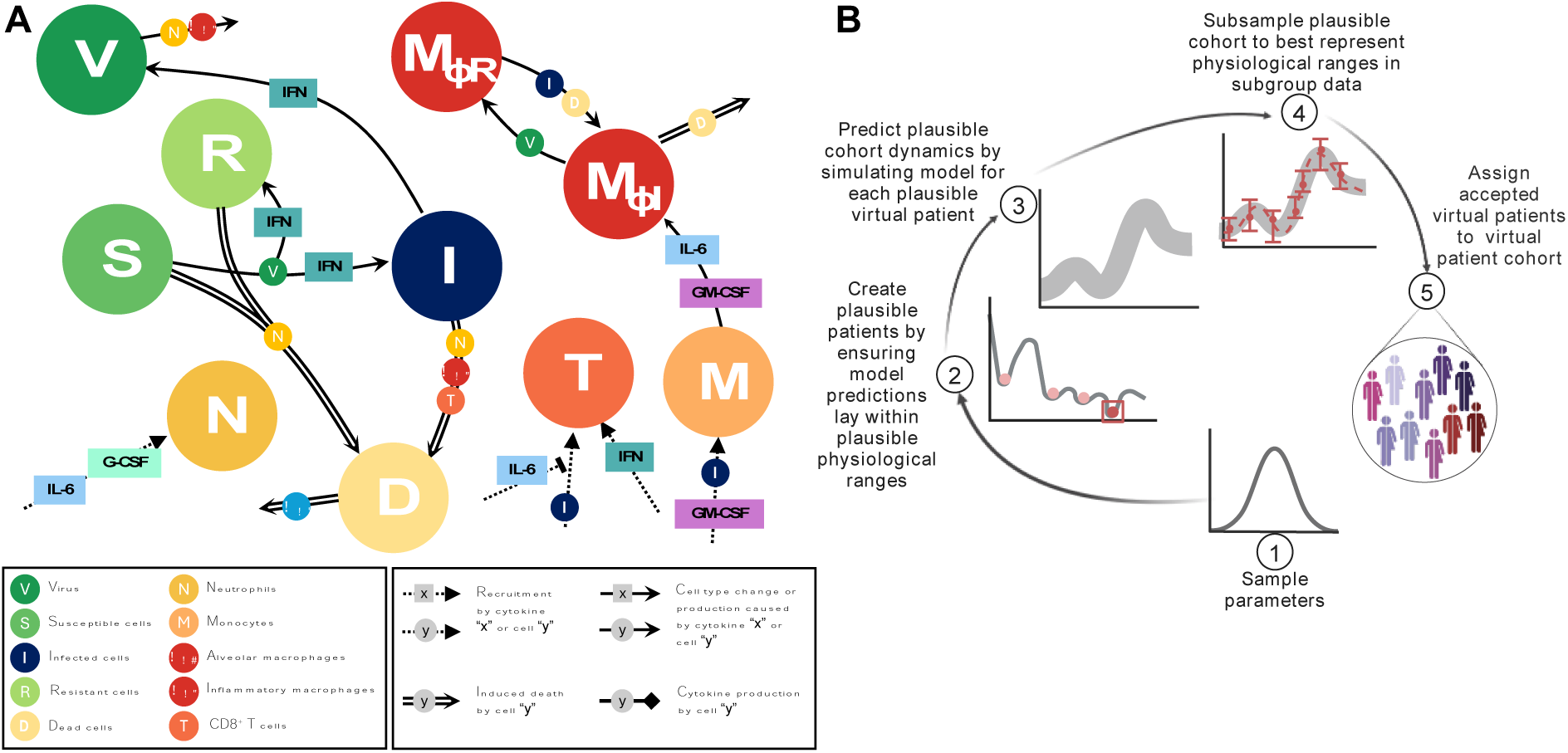
Mathematical model of the systemic immune response to SARS-CoV-2 and virtual patient generation algorithm. A) Mathematical model describing the immune response during COVID-19. Reproduced from Jenner et al.^28^ under CC BY. Virus infects susceptible cells and creates infected or resistant cells based on IFN concentrations. Infected cells die and produce more virus or are eliminated by inflammatory macrophages, neutrophils (recruited by IL-6 and G-CSF), or CD8+ T cells whose population expands based on IFN concentrations and is inhibited by IL-6. Monocytes are recruited by infected cells and differentiate into inflammatory macrophages, which is regulated by GM-CSF and IL-6 concentrations. Some tissue-resident macrophages convert to become inflammatory after encountering infected or dead cells. See Supplementary Information for full model equations and parameter values. B) Schematic description of the virtual patient cohort generation algorithm adapted from Jenner et al.^28^ 1) Parameters associated with macrophage, IL-6, and IFN production are sampled from normal distributions extracted from clinical data (Figure 2). 2) The model is simulated and simulated annealing is performed to minimize the distance between model predictions (outputs) and physiological ranges. 3) Virtual patients whose dynamics fit into the pre-defined ranges are assigned to the cohort of plausible patients. 4) The population of plausible virtual patients is subsampled based on data specific to each studied group (e.g., COVID-19 reference, cancer, immunosuppressed).

In brief, infection begins with virus infecting susceptible lung epithelial cells (*S*), resulting in the production of virus (*V*) and infected cell (*I*) death. Infected cells can secrete IFN-α,β and, depending on the IFN concentration, neighbouring cells may also become resistant (*R*) to viral entry and replication. Infected cells are then removed by cytotoxic CD8+ T cells (*T*), inflammatory macrophages (*M_ΦI_*), and neutrophils (*N*), with neutrophils causing damage to all lung epithelial cells through their release of granules. Monocytes differentiate into inflammatory macrophages based on IL-6 and GM-CSF concentrations, and tissue-resident macrophages (*M_ΦR_*) can transition into inflammatory subsets through contact with either dead or infected cells. Dead cells (*D*) are eliminated by inflammatory macrophages. Neutrophils are recruited by IL-6 and G-CSF, whereas monocytes are attracted by infected cells and GM-CSF. Recruitment of CD8+ T cells is driven by infected cells and IFN and suppressed by IL-6 concentrations (it should be noted that we considered IL-6 as a proxy for the multitude of cytokines that inhibit T cell recruitment). A detailed description of the model equations and parametrization is provided in the Supplementary Information and Jenner et al.^28^. All model simulations were performed in MATLAB^35^ using *ddesd*.

### Generating virtual patient cohorts

To generate the three virtual patient cohorts in our study, we followed the algorithm described in Jenner et al.^28^ (Figure 1B) to ensure that each plausible patient’s immunological trajectory corresponds to available clinical data^4^. The generation process began from the most sensitive parameters (***p***) revealed in the sensitivity analysis in Jenner et al.^28^, including parameters associated with IFN, macrophage and IL-6 production. These were first sampled from normal distributions obtained from available clinical data^30^, with mean values and standard deviations taken from measurements on specific days (see Figure 2). For each virtual patient, the model was simulated and the cost function^36^,

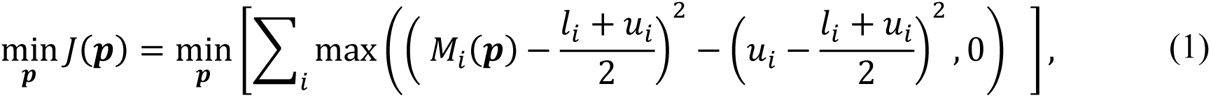

was minimized using simulated annealing. Here, *M_i_*(***p***) is the model output, and *l_i_* and *u_i_* are the lower and upper bounds of each immune population, respectively. We used simulated annealing via the *simulannealbnd* function in Matlab^29^ for this optimization. If predicted dynamics fell within the established data ranges, a virtual patient was accepted as a plausible patient and placed into their respective cohort.

**Figure 2.**
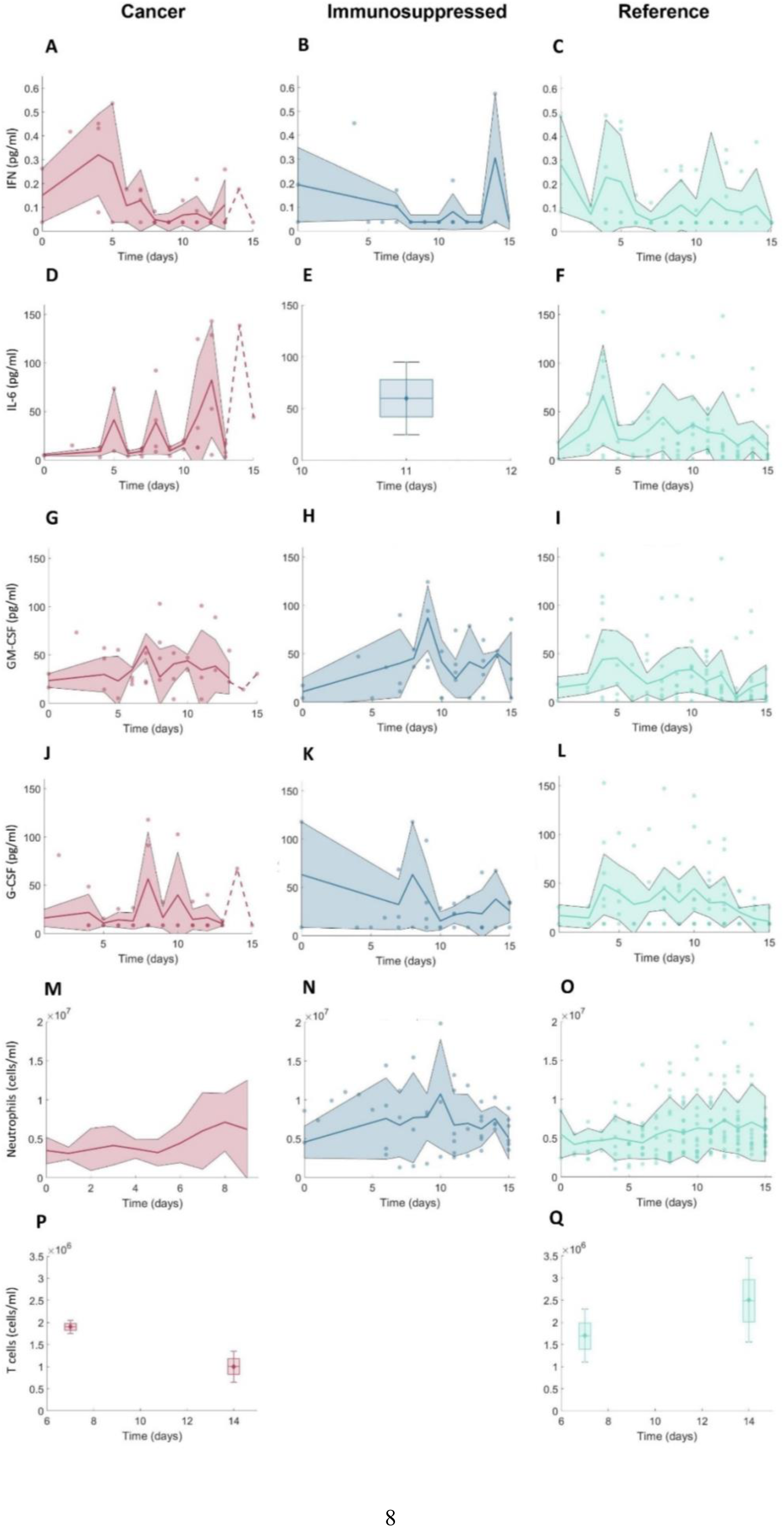
Clinical data measurements from 15 days post-symptom onset in COVID-19+ cancer patients, COVID-19+ immunosuppressed patients, and COVID-19+ patients without cancer or immunosuppression. A-C) IFN concentrations; D-F) IL-6 concentrations; G-I) GM-CSF concentrations; J-L) G-CSF concentrations; M-O) neutrophil concentrations; P-Q) CD8+ T cell concentrations. Note that there were no available CD8+ T cell data from immunosuppressed patients. For further descriptions of the clinical data used in our study, see the Methods and Supplementary Information. Solid lines: mean values. Shaded areas: standard deviations. Dashed lines indicate single observations. Boxplots were used where data were available only on certain days with mean values indicated by diamonds and standard deviations marked by error bars.

Using this approach, we created three cohorts representative of hospitalized and outpatients: 1) COVID-19 cancer, 2) COVID-19 immunosuppressed, and 3) COVID-19 reference (which included patients without cancer or immunosuppression). We then subsampled within each cohort to more tightly match available clinical^29–31^ and reference data^4,5^ (Figure 2; data descriptions can be found in the Supplementary Information). Because COVID-19-positive cancer and immunosuppressed patients tend to have fewer lymphocytes^3,5^ and increased IL-6^4,5^ concentrations, we subsampled virtual patients according to the data from each of these patient groups. For the cancer virtual patient cohort (VPC), we used data from Cai et al.^5^ and clinical data from Montréal hospitals^29–31^ for CD8+ T cells and IL-6 concentrations (Figure 2).

To replicate the neutropenia experienced by patients undergoing chemotherapy^37^, we decreased the initial concentrations of neutrophils for virtual patients in the cancer cohort. For this, we digitized the data from neutropenic patients during SARS-CoV-2 infection described in Lee et al.^37^ using PlotDigitizer^38^. To generate immunosuppressed virtual patients, we used IL-6 concentrations from Monreal et al.^4^. Given their overall higher IL-6 concentrations, these virtual patients also experienced lower CD8+ T cell counts compared to those in the COVID-19 reference group, consistent with our model findings^28^. We assumed that these decreased values were representative of the CD8+ T cell dynamics in hospitalized patients, as CD8+ T cells are lower in immunosuppressed patients due to ongoing treatments. In all, this process resulted in the creation of 280 patients in each of the three cohorts.

### Evaluating disease severity across cohorts using an updated inflammation marker

To compare patient responses across cohorts, we modified the equation for the inflammation marker (Ψ^j^) introduced by Jenner et al.^28^ to evaluate the severity of COVID-19+ virtual patients. This inflammation marker measures each virtual patient’s maximum IL-6 and neutrophil concentrations, and maximal lung tissue damage (i.e., concentration of *D*, see Supplementary Information) according to the mean in the virtual patient cohort. These patient attributes were chosen as they are known to be strongly associated with the final disease outcomes (i.e., disease severity). By comparing patient immune populations to the inflammation marker, our prior work found that IFN peaks were correlated with severity, and that the IFN peak delay defined a severity threshold for Ψ^j^that separated mild from severe cases. Thus, it was concluded that patients with delayed IFN peaks (i.e., those with inflammation marker values above 3) experienced worse (more severe) outcomes.

In our previous work, patient severity was classified based on Ψ^j^ values, but it was done within a single cohort. However, to compare patient responses *between* cohorts, adjustments to the inflammation marker equation introduced in Jenner et al.^28^ were necessary, given that the normalization (denominator terms) in the original equation are specific to the cohort being considered and that these values will vary across cohorts. Thus, we opted here to use the COVID-19 reference cohort as a baseline to measure severity across groups. Accordingly, we modified the denominator values to reflect mean biomarker values from the reference group:

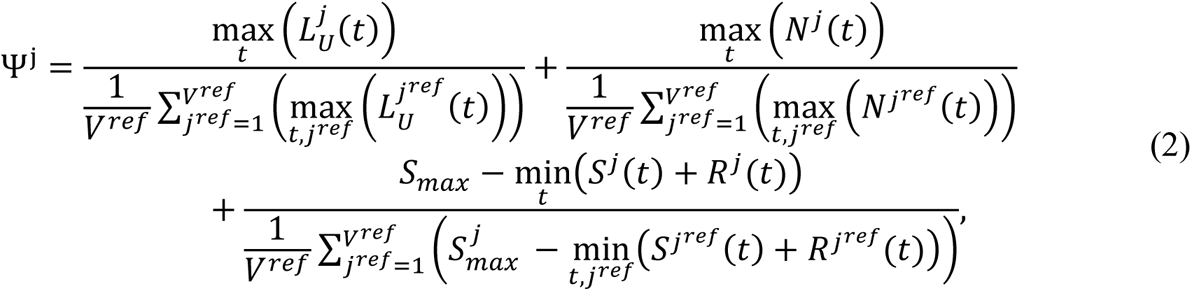

where the index *j* corresponds to the *j*-th patient, *L_U_^j^*, *N^j^*, *S^j^* + *R^j^* and *S_max_* represent the concentrations of unbound IL-6, neutrophils, susceptible plus resistant epithelial cells (undamaged tissue), respectively, and *j^ref^* is the *j*th VP in the COVID-19 reference cohort with *V^ref^* =280 being the total number of patients in the cohort. The change from our previous work allows for inter-cohort comparisons of patient responses, which is crucial as some patients in the vulnerable population virtual cohorts had lower Ψ^j^ values compared to COVID-19 reference patients while at the same time exhibiting markers of increased disease severity (e.g., higher IL-6, lower T cells, etc.) with respect to their own cohort but not necessarily to the others.

### Sensitivity analysis

Given the neutropenic status of the cancer virtual patients in our study, in addition to the sensitivity analysis of full model parameters previously performed in Jenner et al.^28^, we ran a local sensitivity analysis to see how changes in the initial concentration of neutrophils (*N*_0_) may impact other populations and, thus, severity. For this, we varied the initial concentration of neutrophils from 60% to 140% of its baseline value and checked the differences in the output values of certain immune populations (maximal viral concentration, minimum tissue concentration, maximum IFN exposure and maximum concentration of dead cells, inflammatory macrophages, CD8+ effector T cells, IL-6, IFN) compared to their baseline output values. Parameter changes that caused changes greater than 40% were considered significant.

### Statistical analysis

We used the Kolmogorov-Smirnov test at a level of significance of α=0.05 via the *kstest2* function in Matlab^35^ to evaluate statistically significant differences in pair-wise distributions of virtual patient parameters between severe and mild patients across, and among cohorts. To analyze statistical differences in maximal biomarker values observed in virtual patients, we performed ANOVA tests at a level of significance of α=0.05 using *anova1* function in Matlab^35^. To further analyze the statistical differences in parameter values and some maximal biomarker values (e.g., neutrophils, damaged tissue, and IFN concentrations) between certain groups of patients (i.e., severe vs. mild), we performed a pairwise non-parametric Wilcoxon test using the *stat_compare_means* function in R^39^. We considered coefficients of *R* ≥ 0.6 to indicate moderate to strong correlations.

## RESULTS

### Data suggests differences in immune biomarkers values in cancer and immunosuppressed patients compared to those without comorbidities

To ensure all virtual patients trajectories matched the clinical data, we subsampled them based on the biomarker measurements from clinical^29–31^ and reference data^4,5^ sources (Figure 2; see Generating virtual patients cohorts in the Methods and the Supplementary Information). In these data, cancer patients with COVID-19 tended to have decreased T cells count throughout the course of infection compared to cancer-free (reference) individuals with COVID-19 (Figure 2P-2Q). Mean concentrations of IL-6 were increased in both COVID-19+ cancer and immunosuppressed patients groups compared to comorbidity-free group after day 10 post symptom-onset, reaching above 60 pg/ml. The initial value of neutrophils was lowest in COVID-19+ cancer patients, but neutrophil concentrations were similar across the three groups throughout the course of infection. Mean values of GM-CSF were highest in the COVID-19+ immunosuppressed patients (Figure 2G-2I), whereas IFN values tended to decrease in cancer patients during infection, while they remained generally on a constant level in the reference cohort versus the cancer and immunosuppressed patients (Figure 2A-2C).

### Key differences in immunological dynamics of virtual patients in the reference, cancer, and immunosuppressed cohorts

To uncover the potential causes of COVID-19 associated severity in cancer and immunosuppressed populations, we generated virtual patient cohorts each consisting of 280 virtual individuals with COVID-19 who were otherwise healthy (“reference”), had cancer, or were immunosuppressed. The virtual patient selection process (Figure 1B, see Methods) resulted in diverse dynamics. Namely, the cancer (Figure 3A) and immunosuppressed virtual patients (VPs) (Figure 3B) both exhibited significantly decreased CD8+ T cell concentrations (p-values < 10^-8^) compared to the VPs from the reference cohort (Figure 3C). Around 10 days after infection (when concentrations peaked), the mean T cell concentration reached 1.3 × 10^6^cells/ml in the cancer cohort, while it was 1.8 × 10^6^ cells/ml in the COVID-19 reference cohort (Table 1). In comparison, VPs from the immunosuppressed cohort had lower maximal CD8+ T cell concentrations, with a mean of 0.9 × 10^6^ cells/ml (Table 1). These patterns also extended to IL-6 concentrations that similarly varied between groups: immunosuppressed patients had the highest maximal mean values of IL-6 of 60 pg/ml (Figure 3H, Table 1), followed by VPs in the cancer cohort (Figure 3G, Table 1), who were predicted to have an average peak value of 40 pg/ml. In contrast, virtual patients in the COVID-19 reference group had the lowest mean IL-6 peak concentrations of 25 pg/ml (Figure 3I, Table 1), which is consistent with reduced severity in otherwise healthy individuals. Statistical differences in maximal IL-6 values between three groups were confirmed by ANOVA (p-values <10^-8^). A similar trend was observed in GM-CSF, where ANOVA confirmed statistically significant differences in maximal GM-CSF values between cohorts (p-values < 10^-8^); COVID-19 immunosuppressed VPs had highest mean GM-CSF maximal concentration (117 pg/ml), which was almost two times higher than in the COVID-19 reference group (60.17 pg/ml). By comparing maximal values of inflammatory macrophages, we also found statistical differences (p-value < 10^-8^) between all three cohorts. Maximal values of neutrophils were significantly decreased (p-value < 10^-8^) in the cancer cohort compared to other two cohorts (Figure 3J-3L, Table 1). Despite these dissimilarities, our model did not predict a statistically significant difference in the maximal IFN, G-CSF and monocyte concentrations between the three groups (Figure 3D-3F, Table 1).

**Figure 3.**
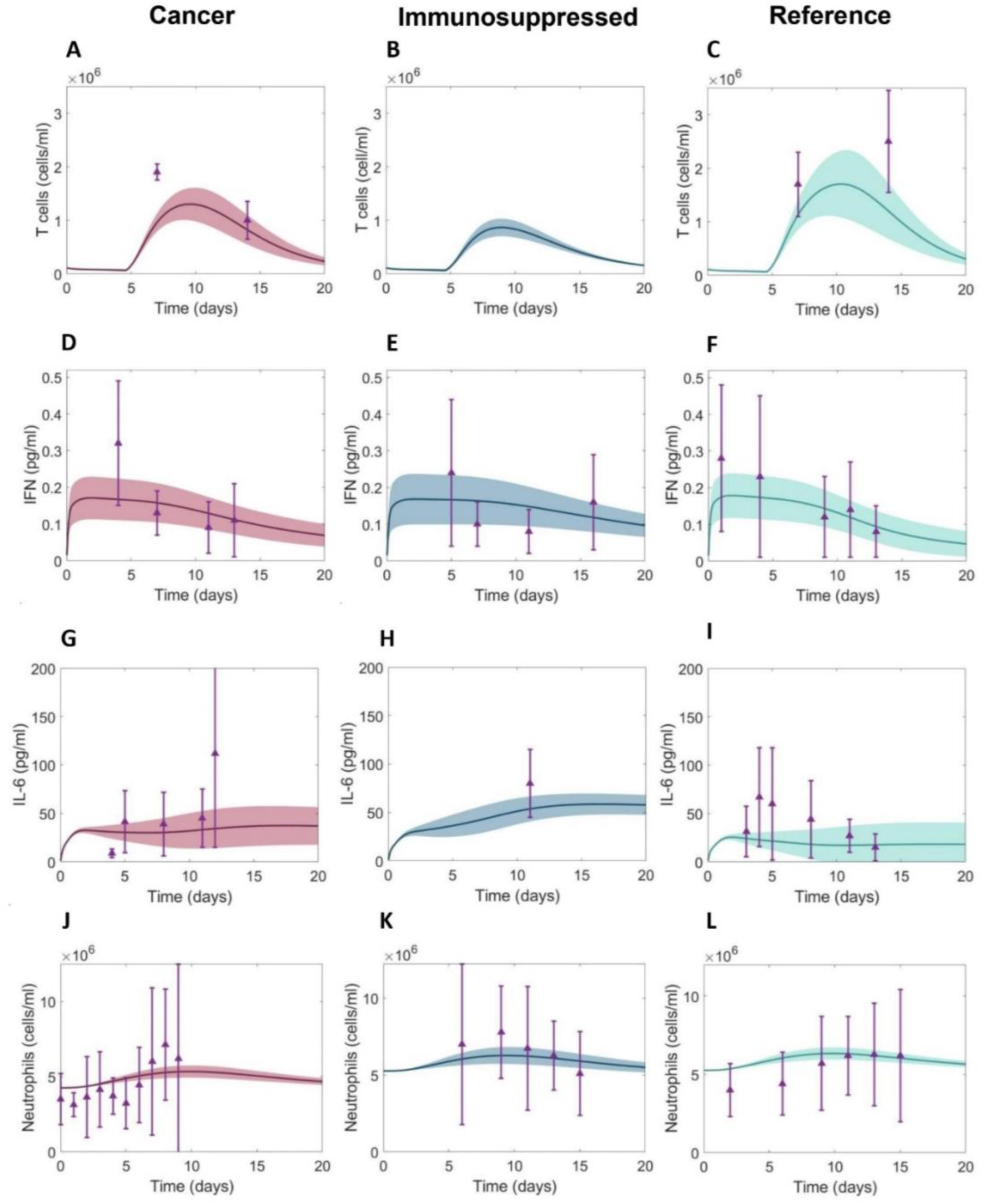
Comparison of immune dynamics in virtual patients from COVID-19+ cancer, immunosuppressed, reference cohorts and clinical data. A-C) CD8+ T cells dynamics, D-F) IFN dynamics, G-I) IL-6 dynamics, J-L) Neutrophils dynamics. Solid curves: mean values within each cohort. Shaded areas: standard deviations. Purple triangles: mean clinical values. Purple vertical lines: standard deviations from clinical observations (Figure 2 and Supplementary Information).

**Table 1.** Predicted peak values over 20 days after infection for the three virtual patient cohorts. Immunosuppressed VPs were found to have the highest maximal inflammatory macrophage, IL-6, and GM-CSF concentrations in addition to the lowest maximal T cell concentrations. In comparison to the COVID-19 reference group, cancer virtual patients were predicted to have higher peak IL-6 and GM-CSF concentrations, increased inflammatory macrophages, and decreased maximal T cells and neutrophils. Values indicate means and standard deviations (SD). * indicates a statistically significant difference (ANOVA) in maximal biomarker values found in patients from cancer and immunosuppressed cohorts versus reference cohort. Statistically significant differences found in maximal biomarker values in patients from cancer versus immunosuppressed cohorts are marked by *.

|  | Units | COVID-19+<br>cancer<br>mean (SD) | COVID-19+<br>immunosuppressed<br>mean (SD) | COVID-19<br>reference<br>mean (SD) |
| --- | --- | --- | --- | --- |
| **T cells** | 10<sup>6</sup> cells/ml | 1.32 (0.28) ** | 0.87 (0.17) ** | 1.78 (0.58) |
| **IFN** | pg/ml | 0.17 (0.06) | 0.17 (0.07) | 0.18 (0.06) |
| **IL-6** | pg/ml | 39.65 (12.95) ** | 58.75 (11.01) ** | 25.15 (12.25) |
| **Neutrophils** | 10<sup>6</sup> cells/ml | 5.35 (0.39) ** | 6.29 (0.52) * | 6.33 (0.36) |
| **GM-CSF** | pg/ml | 78.92 (25.81) ** | 117.01 (21.89) ** | 60.17 (24.43) |
| **G-CSF** | pg/ml | 27.27 (0.83) | 27.18 (1.09) | 27.25 (0.76) |
| **Monocytes** | 10<sup>5</sup> cells/ml | 4.59 (0.21) | 4.59 (0.29) | 4.58 (0.20) |
| **Inflammatory**<br>**macrophages** | 10<sup>5</sup> cells/ml | 1.83 (1.52) ** | 4.38 (2.72) ** | 1.51 (1.53) |

### High tissue damage and increased occurrence of IFN peak delay characterize severe COVID-19 immunosuppressed virtual patients

To uncover mechanistic differences in immune responses in mild and severe COVID-19 virtual patients from vulnerable populations, we compared characteristics (e.g., maximal T cell concentrations) across our three virtual patient cohorts using our updated inflammation marker (Eq. (2)). In all three cohorts, severe patients (patients with high values of severity marker Ψ^j^) tended to have depleted CD8+ T cells (Figure 4A-4C) with the strongest negative correlation (R = -0.88, p-value < 10^-8^) with the inflammation marker (Ψ^j^) found in the cancer cohort. A strong positive correlation (R > 0.9, p-value < 10^-8^) was observed between inflammation marker (Ψ^j^) and maximal IL-6 concentrations (Supplementary Figure 1A-1C), and the maximal concentration of inflammatory macrophages (R > 0.85, p-value < 10^-8^; Supplementary Figure 1D-1F). In both the cancer and immunosuppressed VPCs, we also found a statistically significant weak correlation between severity and peak neutrophils concentrations (R ≈ 0.4, p-value < 10^-8^) in addition to the degree of lung tissue damage (R ≈ 0.5, p-value < 10^-8^) while in the COVID-19 reference cohort, no such relationships were established (maximum neutrophils: R = 0.064, p-value = 0.288; maximum damaged lung tissue: R = -0.108, p-value = 0.072; see Supplementary Figure 1G-1I and 1J-1L). Moderate correlations (R > 0.6, p-value < 10^-8^) between the inflammation marker (Ψ^j^) and peak IFN concentrations were found in both the immunosuppressed and COVID-19 reference cohorts (Figure 4E-4F), with severe immunosuppressed patients (Ψ^j^ > 4) having IFN peak delays more often than patients from other cohorts. In the cancer cohort, we only observed a statistically significant but weak correlation (R < 0.6, p-value < 10^-8^) between IFN peak and Ψ^j^. Together, these findings suggest increased immunological dysregulation in cancer and immunosuppressed virtual patients.

**Figure 4.**
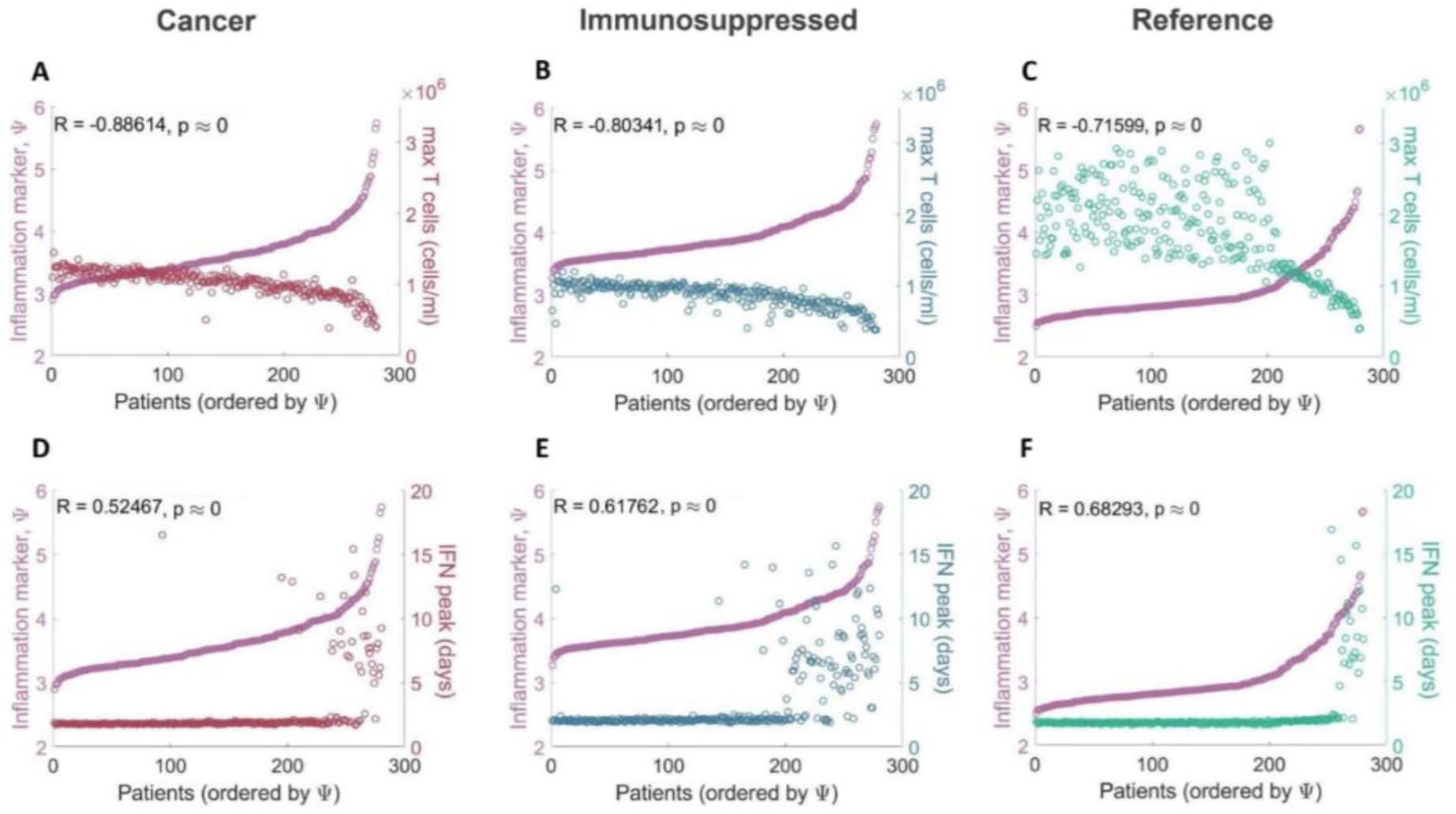
Correlations between maximal T cell and peak IFN concentrations and COVID-19 severity. Maximal T cell concentrations compared to the inflammation marker in patients from the A) COVID-19+ cancer cohort, B) COVID-19+ immunosuppressed cohort, and C) COVID-19+ reference cohort. Maximal T cell concentrations in all three cohorts were found to be negatively correlated with *ψ^j^*. Time to IFN peak concentrations compared to the inflammation marker in patients from the D) COVID-19+ cancer cohort, E) COVID-19+ immunosuppressed cohort, and F) COVID-19+ reference cohort. IFN peak times were positively correlated with *ψ^j^*in the immunosuppressed and COVID-19 reference cohorts. Patients were ordered by *ψ^j^* values, with the mildest patients having the lowest *ψ^j^*values and the most severe the highest *ψ^j^* values.

### Increased monocyte recruitment rates indicate more innate immune dysregulation in the vulnerable groups versus reference patients

We further explored how differences in parameter values led to altered dynamics by selecting virtual patients representing the lowest and highest 10% of inflammation marker values in each VPC and comparing their parameter values. These virtual patients correspond to the mildest or most severe SARS-CoV-2 infections, respectively. We performed statistical analyses using a pairwise non-parametric Wilcoxon test and found increases (p-value < 10^-8^) in the mean values of parameters associated with monocyte-to-macrophage differentiation by IL-6 (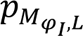; Figure 5A) and decreases (p-value < 0.05) in IFN production rates by infected cells (*p_F,I_*; Figure 5B) in severe patients in all three cohorts. In particular, the values of *p_M,I_* (monocyte recruitment by infected cells; Figure 5C) and *∊_F,I_* (cell-related half-maximal inhibitory (IC50) concentration of IFN on the virus production; Figure 5D) were elevated only in the cancer and immunosuppressed virtual patients with severe COVID-19 (p-value < 0.05); differences between mild and severe virtual patients in the reference cohort were not observed. Values of 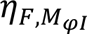 (half-maximal stimulatory (EC50) concentration of inflammatory macrophages on the IFN production; Figure 5E) were increased in cancer and immunosuppressed severe patients, but only in the latter group was the difference statistically significant (p-value < 0.05).

**Figure 5.**
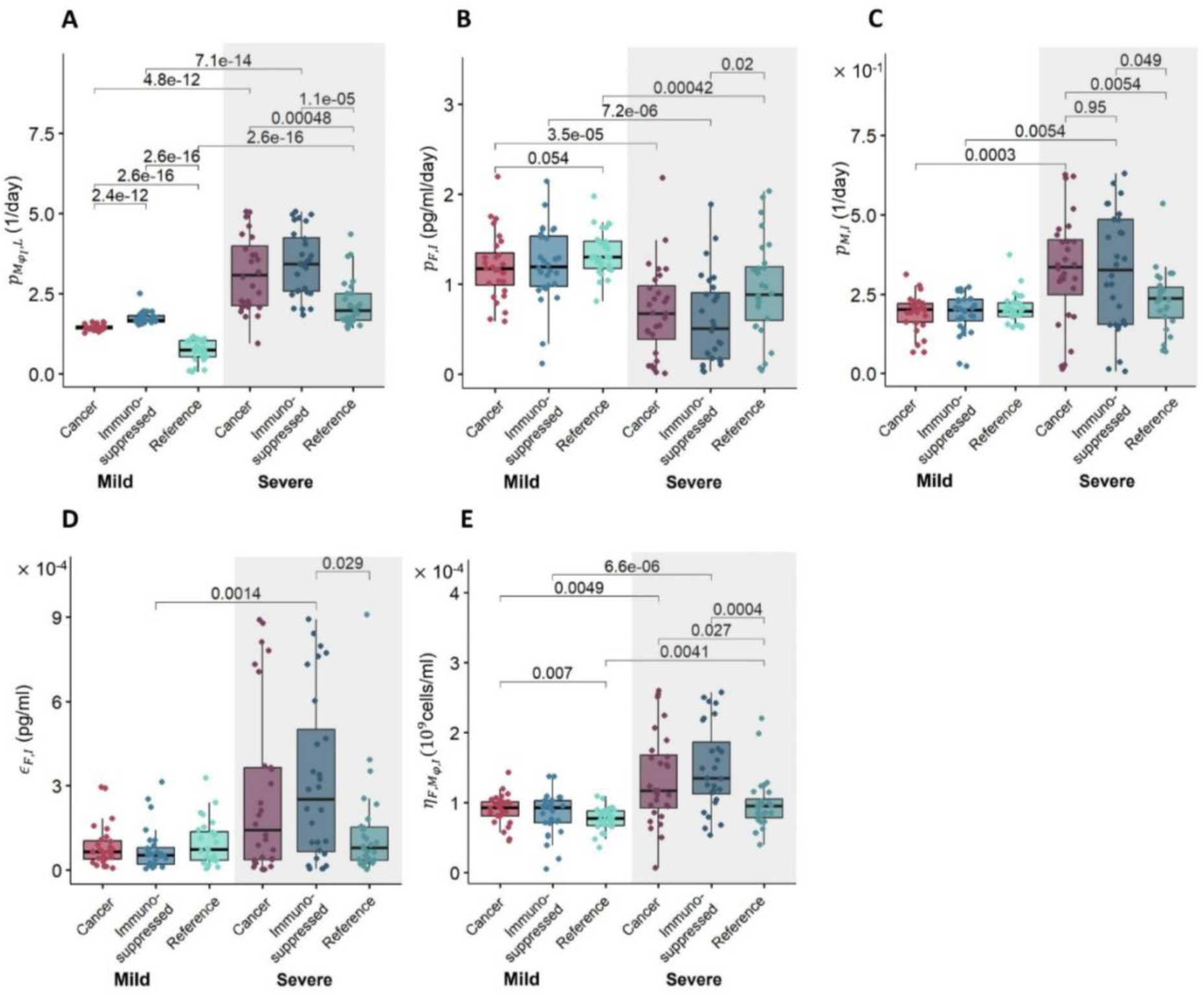
Differences in parameter values between mild and severe patients in each virtual patient cohort. Differences in parameters associated with A) monocyte-to-macrophage differentiation by IL-6 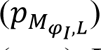, B) IFN production rate by infected cells (*p_F,I_*), C) monocyte recruitment by infected cells (*p_M,I_*), D) cell-related IC50 concentration of IFN on virus production (*∊_F,I_*), and E) EC50 concentration of inflammatory macrophages on the IFN production 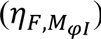. Box plots show the mean values of parameters in each cohort. Statistically significant differences in parameters between groups are marked by p-values above the box plots. A pairwise non-parametric Wilcoxon test was used to assess statistical significance (see Methods).

We then performed a Kolmogorov-Smirnov test to check for statistically significant differences in parameter distributions between the mildest 10% and most severe 10% of virtual patients in all three VPCs (Supplementary Figures 2-9). Three parameters from the cancer VPC were found to differ from the reference VPC when considering the severe virtual patients (Figure 6A-6C). These included the rate of monocyte-to-macrophage differentiation by IL-6 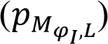, the rate of monocyte recruitment by infected cells (*p_M,I_*), and the IC50 concentration of IFN on virus production (*∊_F,I_*). We also found statistically different distributions of six parameters between the severe immunosuppressed and COVID-19+ reference VPs (Figure 6D-6I), again supporting the observation of increased immune dysregulation in these vulnerable populations. To further characterize the degree of these immunological differences, we also compared parameter distributions between the cancer and immunosuppressed VPCs and found a statistically significant difference in only one parameter (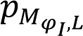 , the monocyte-to-macrophage differentiation by IL-6) between mild patients (Supplementary Figure 8) and no statistical differences between severe patients in those two groups (Supplementary Figure 9).

**Figure 6.**
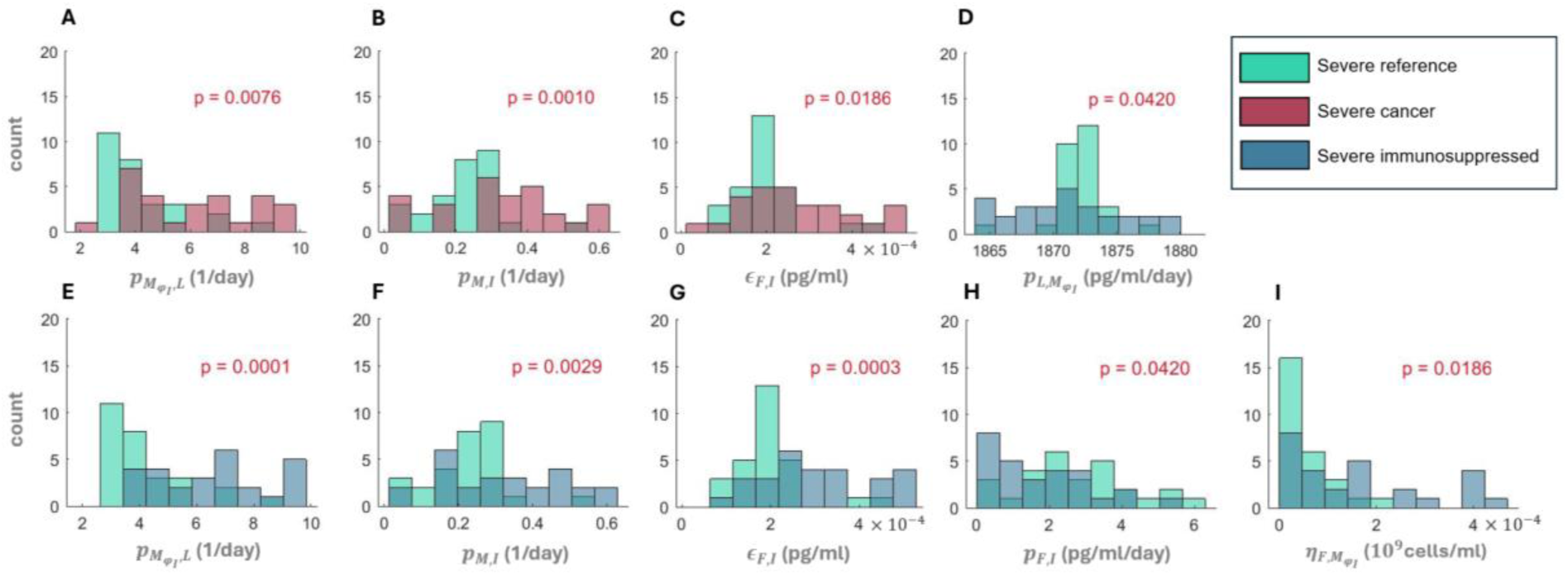
Statistical differences in parameter distributions between severe virtual patients. Statistically significant parameter distributions were evaluated by comparison to the COVID-19 reference cohort using a Kolmogorov-Smirnov test at a level of significance of *α*=0.05. Comparison of the COVID-19 reference to A)-C) severe cancer and D)-I) severe immunosuppressed. The parameter being compared is denoted on the horizontal axis. A) and E) 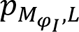 (monocyte-to-macrophage differentiation by IL-6). B) and F) *p_M,I_* (monocyte recruitment by infected cells). C) and G) *∊_F,I_* (cell-related IC50 concentration of IFN on the virus production). D) 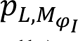 (IL-6 production by inflammatory macrophages). H) *p_F,I_* (IFN production rates by infected cells). I) 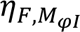 (EC50 concentration of inflammatory macrophages on the IFN production). Red p-values indicate statistically significant differences between parameter distributions.

### Elevated neutrophils are associated with the highest tissue damage in severe cancer and immunosuppressed patients

Using the adjusted inflammation marker, we also examined the relationship between cytokines, cells, and lung tissue damage to distinguish potential severity drivers within each cohort. Overall, we found no correlation between model variables, except for a negative correlation between maximal IFN and damaged tissue (R ≤ -0.6, p-value < 10^-8^) in the three cohorts (Figure 7A-7C). The highest degree of lung tissue damage (marked by red dots) was predicted in the most severe cancer and immunosuppressed patients (compared to mild ones), but not in the most severe patients in the COVID-19 reference cohort (Figure 7C).

**Figure 7.**
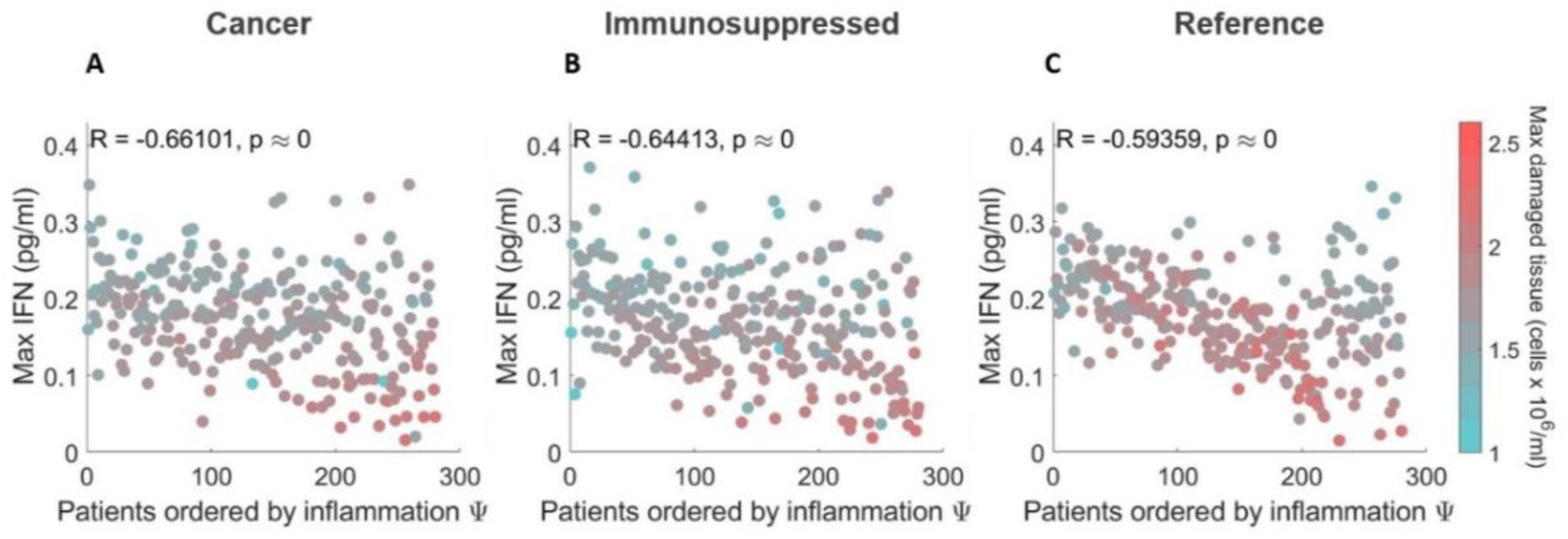
Relationships between maximal IFN and damaged tissue concentrations in virtual patients ordered by severity. A) COVID-19+ patients with cancer. B) COVID-19+ patients with immunosuppression. C) COVID-19+ reference patients. In all three cohorts, maximum IFN concentrations were negatively correlated with the degree of damaged tissue. In the cancer and immunosuppressed cohorts (A and B), the most severe patients (i.e., those with the highest inflammation marker values) were found to have the most tissue damage, contrary to virtual patients in the COVID-19 reference cohort (C). Patients are ordered from the lowest to higest inflammation marker values *ψ^j^*.

To uncover potential factors causing those differences, we analyzed the dynamics and the mean of maximum predicted values of the top 10% of patients (most severe) and bottom 10% (most mild) virtual patients. Our model predicted a comparable amount of damaged tissue over time in the case of severe patients (Supplementary Figure 12). In agreement with findings on lung tissue damage (Figure 7), mean values of maximum damaged tissue were found to be increased only in severe cancer and immunosuppressed patients (Figure 8B).

**Figure 8.**
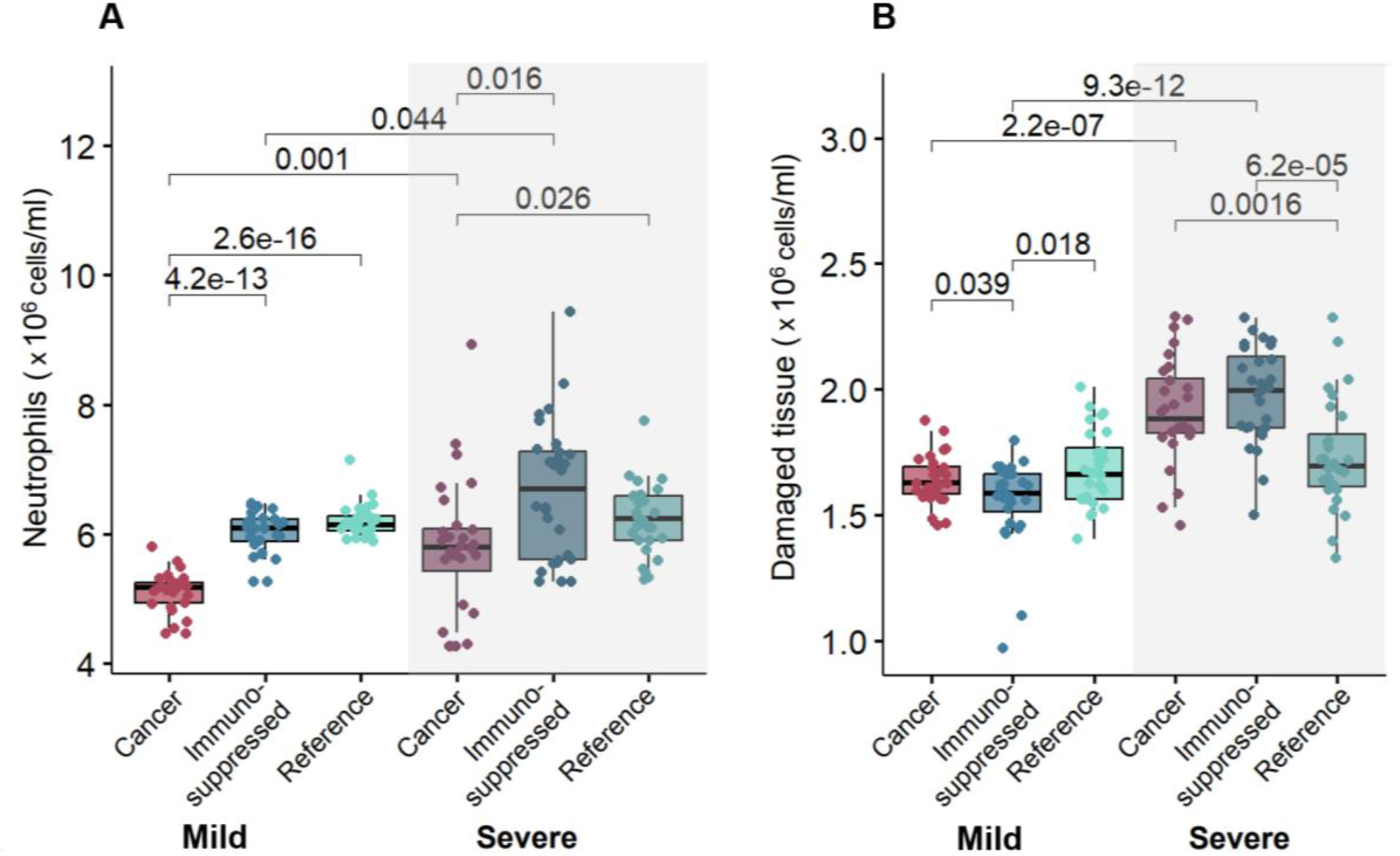
Relationships between maximum neutrophil concentrations and damaged tissue in mild and severe virtual patients. Mean values of A) maximum neutrophils and B) damaged tissue were found to be statistically significantly (p-value < 0.05) increased in cancer and immunosuppressed virtual patients with severe COVID-19 versus those with mild disease. Statistical tests were performed using a pairwise non-parametric Wilcoxon test (see Methods). Statistical differences are marked by p-values above the box plots.

Next, we investigated the dynamics and mean maximum values of other immune populations to look for a potential cause of that feature. Mean values of maximum IFN were decreased in severe patients versus mild patients in all three cohorts (Supplementary Figure 13B). Further, we found that neutrophils were highest in severe immunosuppressed and cancer patients, with this trend also observed in maximum tissue damage (increased values in severe cancer and immunosuppressed patients, Figure 8B) with respect to neutrophil concentrations (Figure 8A). To confirm this, we performed a pairwise non-parametric Wilcoxon test to check for statistical differences. Indeed, we found statistically significant differences in maximal damaged tissue and neutrophils values between mild and severe patients in the cancer and immunosuppressed cohorts (Figure 8), contrary to the reference cohort. While checking for differences in the maximum IFN concentrations, statistical tests confirmed the differences between mild and severe patients in all three cohorts (Supplementary Figure 13B).

### Cancer virtual patients experience overall higher viral loads

Finally, our model predicted higher peak viral loads in both severe and mild cancer virtual patients as compared to virtual patients in the two other cohorts (Figure 9A). We hypothesized that this result was related to depressed initial neutrophil counts (*N*_0_) in these virtual patients caused by chemotherapy-induced neutropenia^40^. To test this, we performed a sensitivity analysis by varying the initial concentration of neutrophils between 60% and 140% of its baseline value (see Methods). Decreasing the initial concentration of neutrophils (*N*_0_) resulted in higher viral loads and maximum IL-6 and IFN concentrations (Figure 9B), seemingly confirming the assumed relationship between initial neutrophil concentrations and viral loads.

**Figure 9.**
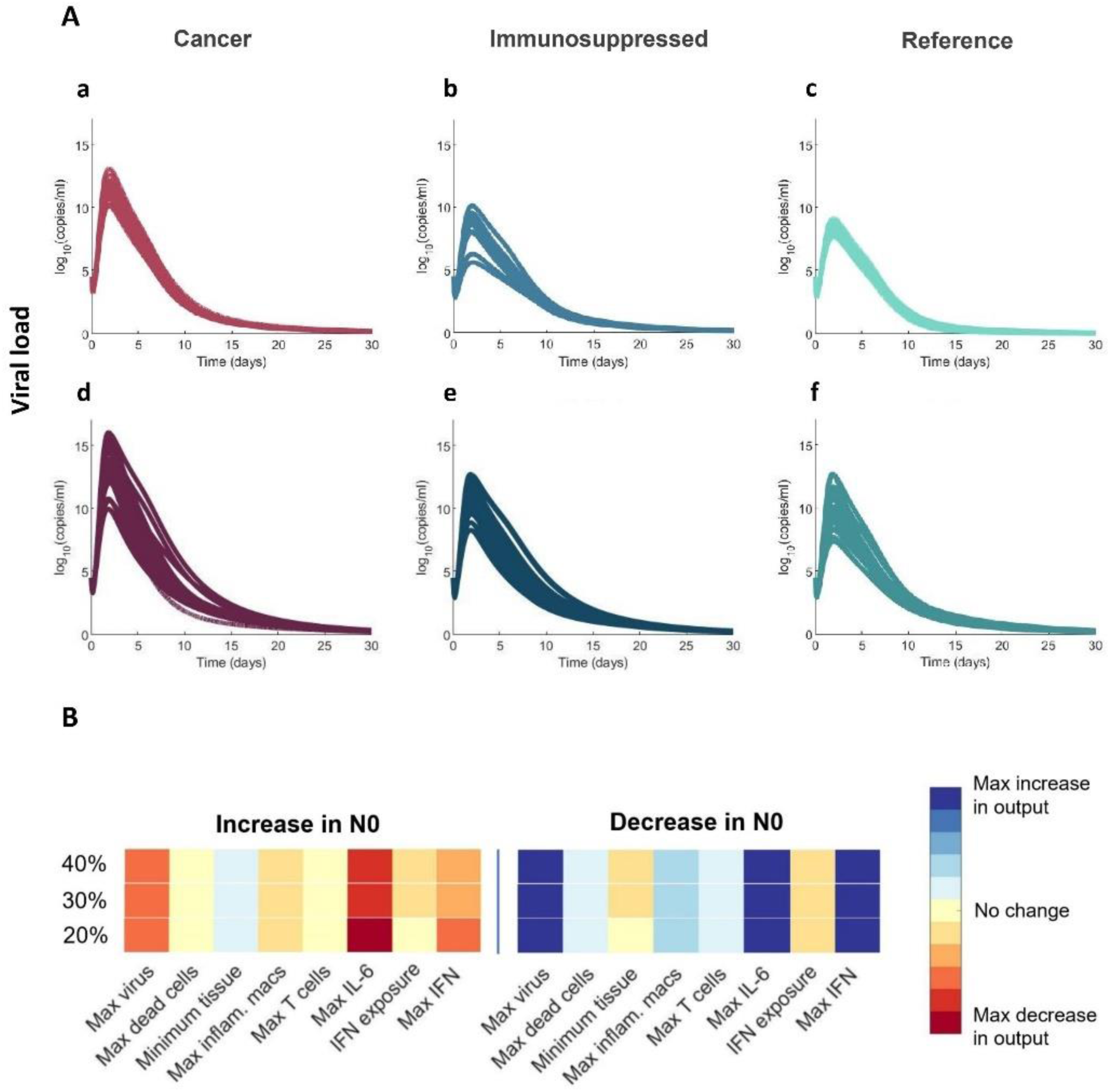
Cancer virtual patients with decreased initial neutrophil concentrations have higher viral load peaks. A) Viral loads in mild a)-c) and severe virtual patients d)-f). B) Sensitivity analysis determined that decreasing *N*_0_(the initial concentration of neutrophils) causes higher peak viral load and peak IL-6 and IFN concentrations.

This suggests that viral load is not the sole driver of severity, which is rather determined through a combination of immunological features^41^.

## DISCUSSION

A better understanding of the immune dynamics and potential causes of severe COVID-19 outcomes in vulnerable groups is essential to improving our understanding of factors driving immune responses during COVID-19, helping to lessen morbidity and mortality in these patients and to select best treatment courses. For example, patients with active cancers are more likely to experience worse COVID-19 disease outcomes as compared to those without cancer^42,43^. Similarly, individuals with immunosuppression triggered by lymphocyte-targeting therapies after organ transplantation have been reported to suffer from severe COVID-19 more often than immunosuppression-free patients^21^. Here, we used mathematical modelling and virtual patient cohorts to predict cellular immune response dynamics in COVID-19 in individuals with cancer or immunosuppression. Based on our previously developed mathematical model^28^, we generated virtual patients whose immunological trajectories corresponded to clinical observations^4,5^. By comparing predicted outcomes between virtual patients with the lowest and highest inflammation marker values both between and within cohorts, we distinguished biomarkers of immune dysregulation and severity, which has implications for drug development and clinical practices.

Our findings suggest that all severe COVID-19 patients, regardless of existing immunosuppression or cancer diagnoses, experience CD8+ T cell depletion, higher IL-6 concentrations, and importantly, delayed type I IFN peaks. Thus, these results further support the role of type I interferons in the control of SARS-CoV-2 infection severity^44^. We also observed delayed IFN peaks in some mild cancer and immunosuppressed virtual patients (Supplementary Figure 10 and 11). Relatedly, previous studies have found that IFN deficiency may be treated by anti-inflammatory therapies that target IL-6^45^. Our model’s predictions further underline the major role of IL-6, which was found to be increased even in mild virtual patients in the cancer and immunosuppressed VPCs.

By comparing parameter values between the top and bottom 10% of virtual patients according to severity (i.e., severe versus mild), we found significant differences in five of the seven parameters used to generate the VPCs. Three of them (monocyte recruitment by infected cells, half-maximal stimulatory concentration of IFN production by inflammatory macrophages, and half-maximal inhibitory concentration of IFN on the virus production) were noticeably increased in severe cancer and immunosuppressed patients (Figure 5), suggesting their roles as potential severity indicators in those groups. Comparing the remaining two parameter values according to severity revealed differences between and within cohorts. For example, the rate of monocyte-to-macrophage differentiation by IL-6 tended to be increased in severe patients and was highest in the immunosuppressed and cancer cohorts and lowest in the COVID-19 reference cohort. This agrees with clinical findings from circulating blood cells those in the lungs in severe COVID-19 patients^46^. Further, IL-6 concentrations are higher in immunosuppressed patients^4^, in agreement with our model predictions. The rate of IFN production by infected cells was also predicted to be highest in mild patients in the COVID-19 reference cohort as compared to the other two cohorts, again showing the significant role of IFN in coordinating a sufficient immune defense against SARS-CoV-2 infection.

Although neutrophils play a crucial role in blocking fungal and bacterial infections^47^, their function in viral infection is not yet fully establlished^48^. When we decreased the initial neutrophil concentration to mimic neutropenia characteristic of cancer patients, our model predicted higher peak viral loads in both severe and mild virtual cancer patients (Figure 9A). In our model, neutrophils quickly remove free viral particles and cause damage to all cells (including infected cells), hence a lower initial concentration of these cells may result in a higher number of infected cells leading to increased viral production at the beginning of infection. Later, when neutrophil concentrations in cancer virtual patients were predicted to reach comparable levels to virtual patients in the other cohorts (around day 7 post-infection), we observed a similar trend in the viral load dynamics, namely that its concentration also decreased and was comparable to those predicted in the two other cohorts. As a higher viral load peak was observed in severe patients, our results suggest that pre-existing neutropenia in cancer patients may be associated with adverse outcomes, consistent with findings from other studies^15^. However, throughout the course of infection, our model predicts that cancer patients with severe COVID-19 will nonetheless experience neutrophilia (Figure 8A), in agreement with clinical studies like that of Lee et al.^37^.

Overall, our study supports the continued investigation of longitudinal immunological dynamics in groups vulnerable to COVID-19 by highlighting key mechanistic differences in their immune responses. In particular, our results revealed the effect of pre-existing neutropenia on viral load in cancer patients, which can result in a more severe course of infection^49^. However, we also found an association between elevated neutrophils and high tissue damage in severe COVID-19+ cancer and immunosuppressed patients, which suggests the potential danger of neutrophilia even during immunosuppression. This may explain why some studies found no connection between neutropenia and severe COVID-19^37^, and even found decreased neutrophils beneficial. However, as other studies identified low neutrophil count as a potential risk factor^15^ in COVID-19, our findings support accounting for neutropenia in treatment decisions. Interestingly, when considering full cohorts, we did not find any correlations between maximum neutrophil count and damaged tissue. This lack of association is notable given the key roles of cell-mediated immunity during infection with SARS-CoV-2. For example, through the release of neutrophil extracellular traps and reactive oxygen species, neutrophils can cause extensive damage to tissues, so a correlation between maximum neutrophil count and damaged tissue would be expected in all patients. Moreover, by adjusting immunological trajectories to available CD8+ T cell and IL-6 data, our model predicted elevated pro-inflammatory compounds (such as GM-CSF) in both cancer and immunosuppressed patients. Thus, our results suggest the consideration of inhibitory therapies, as GM-CSF has been identified as a driver of lung tissue damage^50^, and underline the delicate balance that must be struck to generate a robust yet controlled response to SARS-CoV-2. Together, this work puts forward a hypothesis for increased severity in both cancer and immunosuppressed patients, whose immunological systems are dysregulated either through disease or by immunomodulatory treatments.

Virtual populations based on mechanistic mathematical models enable the study and prediction of immune responses to viruses or vaccination without the need for extensive amounts of clinical data, making the approach a promising tool to study emerging infectious diseases and a variety of other contexts^28,51–54^. Nonetheless, our model has limitations. Specifically, certain innate immune cells (i.e., natural killer cells^55^) and cytokines that play an important role in fighting SARS-CoV-2 (i.e., IL-1, IL-12, TNF-*α*^56^) were not considered in our model. However, the major model component IL-6, the main driver of T cell depletion throughout the infection, mimics the effects of other cytokines and drugs that inhibits T cell recruitment, suggesting our results can be extended to other anti-inflammatory cytokines not included in our model. Further, our model did not account for the humoral response provided by B cells and antibodies, and thus, it has limited application to vaccination studies. Adding those components would enable the identification of other severity-associated factors and significantly improve our understanding of the intricate dynamics of the immune response to SARS-CoV-2 and other viral infections. Further, the addition of humoral immunity to our model would enable the identification of the mechanisms of decreased vaccine efficacy, as reported in many studies^11,57^, in vulnerable groups.

In summary, our findings corroborate that unregulated immune responses in cancer and immunosuppressed patients place them in a high-risk position of experiencing severe COVID-19. Furthermore, the approach presented here can be used in complement to experimental and clinical studies of COVID-19 and other viral respiratory diseases to comprehensively explore immune response kinetics after infection, thereby improving our understanding of the disease severity.

## ACKNOWLEDGEMENTS

We thank all the patients, family members, and staff from all the units that participated in the study, in addition to the teams of Drs Madeleine Durand, Michaël Chassé, Brent Richards, Daniel Kaufmann, and technicians from other laboratories at the CRCHUM for patient recruitment, sample collection, and blood specimen processing. We acknowledge the clinical research teams for clinical data retrieval, J Plantin, C Dufour, I Turcotte, and R Fromentin for sample collection and processing, and the help of Marc Messier Peet, Pascale Arlotto, Nakome Nguissan, Fatma Mayil, Maya Salame, and Nathalie Brassard for participant recruitment at CHUM.

## CODE AVAILABILITY

The computational code to simulate the full immunological model is available on GitHub at https://github.com/adriannejnner/COVID19-Virtual-Trial-PLOS-Pathogens. Matlab arrays for each of the three virtual patient cohorts are available on GitHub at https://github.com/mlcraig/COVID19-Virtual-Trial-vulnerable-groups.git.

