## Supplementary Information for "Using virtual patient cohorts to uncover immune response differences in cancer and immunosuppressed COVID-19 patients"

In our work, we applied a mathematical model describing the immune response against SARS-CoV-2 from Jenner et al.[1] consisting of 17 differential equations (see below). Model schematics and further descriptions are provided in the Main Text.

### MODEL EQUATIONS

$$\frac{dV}{dt} = pI - \delta_{V,M\Phi}M_{\Phi I}V - \delta_{V,N}NV - d_V V,$$

$$\frac{dS}{dt} = \lambda_S\left(1 - \frac{S+I+R+D}{S_{max}}\right)S - \beta SV - \frac{\rho\delta_N S N^{h_N}}{N^{h_N} + IC_{50,N}^{h_N}},$$

$$\frac{dI}{dt} = \frac{\beta_{\epsilon_{F,I}}}{F_B + \epsilon_{F,I}}S(t-\tau_I)V(t-\tau_I) - d_I I - \frac{\delta_N I N^{h_N}}{N^{h_N} + IC_{50,N}^{h_N}} - \delta_{I,M\Phi}M_{\Phi I}I - \delta_{I,T}TI,$$

$$\frac{dR}{dt} = \lambda_S\left(1 - \frac{S+I+R+D}{S_{max}}\right)R + \frac{\beta F_B}{F_B + \epsilon_{F,I}}S(t-\tau_I)V(t-\tau_I) - \frac{\rho\delta_N R N^{h_N}}{N^{h_N} + IC_{50,N}^{h_N}},$$

$$\frac{dD}{dt} = d_I I + \frac{\delta_N(\rho S + \rho R + I)N^{h_N}}{N^{h_N} + IC_{50,N}^{h_N}} + \delta_{I,M\Phi}(M_{\Phi I})I + \delta_{I,T}TI - d_D D + \left(\delta_{M\Phi,D} - \delta_{D,M\Phi}\right)(M_{\Phi R} + M_{\Phi I})D,$$

$$\frac{dM_{\Phi R}}{dt} = -a_{I,M\Phi}M_{\Phi R}(I+D) - \delta_{M\Phi,\mathrm{D}}DM_{\Phi R} + \left(1 - \frac{M_{\Phi R}}{M_{\Phi max}}\right)\frac{\lambda_{M_\Phi}M_{\Phi I}}{V + \epsilon_{V,M\Phi}} - d_{M_{\Phi R}}M_{\Phi R},$$

$$\frac{dM_{\Phi I}}{dt} = a_{I,M\Phi}M_{\Phi R}(I+D) + \frac{p_{M_{\Phi I},G}G_B^{h_{M,M\Phi}}M}{G_B^{h_{M,M\Phi}} + \epsilon_{G,M_{\Phi I}}^{h_{M,M\Phi}}} + \frac{p_{M_{\Phi I},L}L_B M}{L_B + \epsilon_{L,M_\Phi}} - d_{M_{\Phi I}}M_{\Phi I} - \delta_{M_\Phi,D}DM_{\Phi I} - \left(1 - \frac{M_{\Phi R}}{M_{\Phi max}}\right)\frac{\lambda_{M_\Phi}M_{\Phi I}}{V + \epsilon_{V,M_\Phi}},$$

$$\frac{dM}{dt} = \left(M_{prod}^* + \left(\psi_M^{max} - M_{prod}^*\right)\frac{G_B^{h_M}}{G_B^{h_M} + \epsilon_{G,M}^{h_M}}\right)M_R + \frac{p_{M,I}IM}{I + \epsilon_{I,M}} - \frac{p_{M_{\Phi I},G}G_B^{h_{M,M\Phi}}M}{G_B^{h_{M,M\Phi}} + \epsilon_{G,M_{\Phi I}}^{h_{M,M\Phi}}} - \frac{p_{M_{\Phi I},L}L_B M}{L_B + \epsilon_{L,M_\Phi}} - d_M M,$$

$$\frac{dN}{dt} = \left(N^*_{prod} + \left(\psi_N^{max} - N^*_{prod}\right)\frac{C_{BF} - C^*_{BF}}{C_{BF} - C^*_{BF} + \epsilon_{C,N}}\right)N_R + \frac{p_{N,L}L_B}{L_B + \epsilon_{L,N}} - d_N N,$$

$$\frac{dT}{dt} = \frac{p_{T,I}I(t-\tau_T)\epsilon_{L.T}}{L_B + \epsilon_{L,T}} + \frac{p_{T,F}F_B T}{F_B + \epsilon_{F,T}} - d_T T,$$

$$\frac{dL_U}{dt} = \frac{p_{L.I}I}{I + \eta_{L,I}} + \frac{p_{L,M_{\Phi I}}M_{\Phi I}}{M_{\Phi I} + \eta_{L,M_{\Phi I}}} + \frac{p_{L,M}M}{M + \eta_{L,M}} - k_{lin_L}L_U - k_{B_L}\left((M+N+T)A_L - L_B\right)L_U + k_{U_L}L_B,$$

$$\frac{dL_B}{dt} = -k_{int_L}L_B + k_{B_L}\left((M+N+T)A_L - L_B\right)L_U - k_{U_L}L_B,$$

$$\frac{dG_U}{dt} = \frac{p_{G,M_{\Phi I}}M_{\Phi I}}{M_{\Phi I} + \eta_{G,M_\Phi}} + \frac{p_{G.M}M}{M + \eta_{G,M}} - k_{lin_G}G_U - k_{B_G}(MA_G - G_B)G_U + k_{U_G G_B},$$

$$\frac{dG_B}{dt} = -k_{int_G}G_B + k_{B_G}(MA_G - G_B)G_U - k_{U_G}G_B,$$

$$\frac{dC_U}{dt} = \frac{p_{C,M}M}{M + \eta_{C,M}} - k_{lin_C}C_U - k_{B_C}(NA_C - C_B)(C_U)^{POW} + k_{U_C}C_B,$$

$$\frac{dC_B}{dt} = -k_{int_C}C_B + k_{B_C}(NA_C - C_B)(C_U)^{POW} - k_{U_C}C_B,$$

$$\frac{dF_B}{dt} = -k_{int_F}F_B + k_{B_F}\left((T+I)A_F - F_B\right)F_U - k_{U_F}F_B,$$

where

$$A_L = \frac{MM_L}{6.02214 \times 10^{23}}\left(K_{L,N} + K_{L,T} + K_{L,M}\right) \cdot \left(\frac{10^{-3}}{5000}\right),$$

$$A_G = \frac{MM_G}{6.02214 \times 10^{23}}K_{G,M} \cdot \left(\frac{10^{-3}}{5000}\right),$$

$$A_C = \hat{p}\frac{MM_C}{6.02214 \times 10^{23}}K_{C,N} \cdot \left(\frac{10^{1}}{5000}\right),$$

$$A_F = \frac{MM_F}{6.02214 \times 10^{23}}\left(K_{F,T} + K_{F,I}\right) \cdot \left(\frac{10^{-3}}{5000}\right).$$

### PARAMETER VALUES

| Parameter | Description | Value | Units |
|---|---|---|---|
| $p$ | Lytic viral production rate | 2.59 | 1/day×log(cop/ml)/$10^9$cells |
| $\lambda_S$ | Proliferation of epithelial cells | 0.74 | 1/day |
| $S_{max}$ | Epithelial cells carrying capacity | $S_0$ | $10^9$ cells |
| $\lambda_{M\Phi}$ | Production of alveolar macrophages | 5943 | log(cop/ml)/day |
| $M_{\Phi max}$ | Alveolar macrophage carrying capacity | $M_{\Phi R,0}$ | $10^9$ cells/ml |
| $\beta$ | SARS-CoV-2 virus infection rate | 0.29 | 1/day×1/log(cop/ml) |
| $\tau_I$ | Eclipse time | 0.17 | day |
| $d_I$ | Death rate of infected cells | 0.1 | 1/day |
| $\tau_T$ | Delay in CD8+ T cells arrival | 4.5 | 1/day |

**Supplementary Table 1. Viral kinetic parameters.**

| Parameter | Description | Value | Units |
|---|---|---|---|
| $p_{M_{\Phi I},G}$ | Monocyte-to-macrophage differentiation by GM-CSF | 1.7 | 1/day |
| $p_{M_{\Phi I},L}$ | Monocyte-to-macrophage differentiation by IL-6 | 1.7 | 1/day |
| $a_{I,M\Phi}$ | Activation of macrophages by infected and dead cells | $1.1\times 10^3$ | ml/($10^9$cells)×(1/day) |
| $p_{M,I}$ | Monocyte recruitment rate by infected cells | 0.22 | 1/day |
| $p_{T,F}$ | CD8+ T cell production rate by IFN | 4 | 1/day |
| $p_{N,L}$ | Neutrophils recruitment rate by IL-6 | 0.21 | 1/day |
| $p_{T,L}$ | CD8+ T cell recruitment rate by IL-6 | 4 | 1/day |
| $p_{T,I}$ | CD8+ T cell proliferation rate | 1 | 1/day |
| $M^*_{prod}$ | Homeostasis reservoir release rate | 0.13 | 1/day |
| $\psi_M^{max}$ | Maximal reservoir release rate | 11.55 | 1/day |
| $N^*_{prod}$ | Homeostasis reservoir release rate | 0.21 | 1/day |
| $\psi_N^{max}$ | Maximal reservoir release rate | 4.13 | 1/day |
| $C^*_{BF}$ | Homeostasis neutrophil receptor bound fraction | $1.6\times 10^{-5}$ | Unitless |

**Supplementary Table 2. Cell production, recruitment, and activation rates.**

| Parameter | Description | Value | Units |
|---|---|---|---|
| $\epsilon_{F,I}$ | Cell-related half-maximal inhibitory (IC50) concentration of IFN on the virus production | $4.7\times 10^{-4}$ | pg/ml |
| $\epsilon_{L,M_\Phi}$ | Cell-related IC50 concentration of IL-6 on monocytes to macrophage differentiation | 0.011 | pg/ml |
| $\epsilon_{G,M_{\Phi I}}$ | Cell-related IC50 concentration of GM-CSF on monocyte to macrophage differentiation | 0.027 | pg/ml |
| $\epsilon_{G,M}$ | Cell-related IC50 concentration of GM-CSF recruitment of monocytes | 57.2 | pg/ml |

|  |  |  |  |
|---|---|---|---|
| $\epsilon_{F,T}$ | Cell-related IC50 concentration of IFN production of CD8+ T cells | 0.004 | pg/ml |
| $\epsilon_{C,N}$ | Cell-related IC50 concentration of G-CSF recruitment of neutrophils | $1.89\times 10^{-4}$ | Unitless |
| $\epsilon_{L,N}$ | Cell-related IC50 concentration of IL-6 recruitment of neutrophils | 57.2 | pg/ml |
| $\epsilon_{I,M}$ | Cell-related IC50 concentration of infected cell monocyte recruitment | 0.11 | $10^9$ cells/ml |
| $\epsilon_{L,T}$ | Cell-related IC50 concentration of IL-6 production of CD8+ T cells | $3 \times 10^{-4}$ | pg/ml |
| $\epsilon_{V,M_\Phi}$ | Cell-related IC50 concentration of viral load for mac replenishing | 2.96 | log(cop/ml) |
| $\epsilon_{T,I}$ | Cell-related IC50 effect concentration of antigen-driven proliferation | $10^{-6}$ | $10^9$ cells/ml |
| $h_M$ | Hill coefficient of GM-CSF monocyte recruitment | 1.67 | Unitless |
| $h_{M,M_\Phi}$ | Hill coefficient of GM-CSF monocyte to macrophages | 2.03 | Unitless |
| $h_N$ | Hill coefficient of neutrophil-induced damage | 3.02 | Unitless |
| $IC_{50,N}$ | Neutrophil IC50 concentration of neutrophil-induced damage | 0.047 | $10^9$ cells/ml |

**Supplementary Table 3. Cell-related half-effect ($\epsilon$), IC50 ($IC_{50}$), and Hill coefficient (h) parameters**

| Parameter | Description | Value | Units |
|---|---|---|---|
| $\delta_{V,M\Phi}$ | Rate of viral clearance by macrophages | 768 | ml/($10^9$ cells) $\times$ 1/day |
| $\delta_{V,N}$ | Rate of viral clearance by neutrophils | 2304 | ml/($10^9$ cells) $\times$ 1/day |
| $\delta_N$ | Rate of neutrophil-inflicted damage | 1.68 | 1/day |
| $\rho$ | Bystander death modulation constant | 0.5 | Unitless |
| $\delta_{I,M\Phi}$ | Rate macrophages phagocytose infected cells | 121 | ml/($10^9$ cells) $\times$ 1/day |
| $\delta_{I,T}$ | Rate CD8+ T cells induce apoptosis in infected cells | 238 | ml/($10^9$ cells) $\times$ 1/day |
| $\delta_{M\Phi,\mathrm{D}}$ | Rate macrophages die from phagocytosis | 6.06 | ml/($10^9$ cells) $\times$ 1/day |
| $\delta_{D,M\Phi}$ | Rate macrophages phagocytose dead cells | 8.03 | ml/($10^9$ cells) $\times$ 1/day |

**Supplementary Table 4. Cell- and virus-induced death rates**

| Parameter | Description | Value | Units |
|---|---|---|---|
| $d_V$ | Viral decay rate | 1.81 | 1/day |
| $d_D$ | Degradation rate of apoptotic cells | 8 | 1/day |
| $d_{M_{\Phi\mathrm{R}}}$ | Alveolar macrophage death rate | 0 | 1/day |
| $d_{M_{\Phi\mathrm{I}}}$ | Inflammatory macrophage death rate | 0.3 | 1/day |
| $d_M$ | Monocyte death rate | 0.76 | 1/day |
| $d_N$ | Neutrophil death rate | 1.28 | 1/day |
| $d_T$ | CD8+ T cell death rate | 0.4 | 1/day |

**Supplementary Table 5. Cell death and virus decay rates.**

| Parameter | Description | Value | Units |
|---|---|---|---|
| $p_{L,I}$ | Rate of IL-6 production by infected cells | 11.89 | pg/ml/day |
| $p_{L,M_{\Phi I}}$ | Rate of IL-6 production by inflammatory macrophages | 1872 | pg/ml/day |
| $p_{L,M}$ | Rate of IL-6 production by monocytes | 72.56 | pg/ml/day |
| $p_{G,M_{\Phi I}}$ | Rate of GM-CSF production by inflammatory macrophages | 2626 | pg/ml/day |
| $p_{C,M}$ | Rate of G-CSF production by monocytes | 26.26 | pg/ml/day |
| $p_{G,M}$ | Rate of GM-CSF production by monocytes | 3070 | pg/ml/day |
| $p_{F,I}$ | Rate of IFN production by infected cells | 2.82 | pg/ml/day |
| $p_{F,M_{\Phi I}}$ | Rate of IFN production by inflammatory macrophages | 1.3 | pg/ml/day |
| $p_{F,M}$ | Rate of IFN production by monocytes | 3.56 | pg/ml/day |

**Supplementary Table 6. Cytokine production rates.**

| Parameter | Description | Value | Units |
|---|---|---|---|
| $\eta_{L,I}$ | Half-maximal stimulatory (EC50) concentration of infected cells on the IL-6 production | 0.7 | $10^9$ cells/ml |
| $\eta_{L,M}$ | EC50 concentration of monocytes on the IL-6 production | 0.0045 | $10^9$ cells/ml |
| $\eta_{L,M_{\Phi I}}$ | EC50 concentration of inflammatory macrophages on the IL-6 production | $3.6\times 10^{-5}$ | $10^9$ cells/ml |
| $\eta_{G,M_{\Phi I}}$ | EC50 concentration of inflammatory macrophages on the GM-CSF production | $3.6\times 10^{-5}$ | $10^9$ cells/ml |
| $\eta_{G,M}$ | EC50 concentration of monocytes on the GM-CSF production | 0.15 | $10^9$ cells/ml |
| $\eta_{C,M}$ | EC50 concentration of monocytes on the G-CSF production | 3.05 | $10^9$ cells/ml |
| $\eta_{F,I}$ | Half-effect concentration of IFN production by infected cells | 0.011 | $10^9$ cells/ml |
| $\eta_{F,M_{\Phi I}}$ | EC50 concentration of inflammatory macrophages on the IFN production | $1.3\times 10^{-6}$ | $10^9$ cells/ml |
| $\eta_{F,M}$ | EC50 concentration of monocytes on the IFN production | 0.54 | $10^9$ cells/ml |

**Supplementary Table 7. Cytokine production regulation parameters.**

| Parameter | Description | Value | Units |
|---|---|---|---|
| $k_{lin_L}$ | Rate of IL-6 renal clearance | 16.6 | 1/day |

| | | | |
|---|---|---|---|
| $k_{lin_G}$ | Rate of GM-CSF renal clearance | 11.7 | 1/day |
| $k_{lin_C}$ | Rate of G-CSF renal clearance | 0.16 | 1/day |
| $k_{lin_F}$ | Rate of IFN renal clearance | 18 | 1/day |
| $k_{int_L}$ | Internalization rate of IL-6 | 61.8 | 1/day |
| $k_{int_G}$ | Internalization rate of GM-CSF | 73.4 | 1/day |
| $k_{int_C}$ | Internalization rate of G-CSF | 462 | 1/day |
| $k_{int_F}$ | Internalization rate of IFN | 17 | 1/day |

***Supplementary Table 8. Cytokine linear (renal) clearance and internalization rates.***

| Parameter | Description | Value | Units |
|---|---|---|---|
| $k_{B_L}$ | IL-6 binding rate | 0.0018 | ml/pg/day |
| $k_{B_G}$ | GM-CSF binding rate | 0.0021 | ml/pg/day |
| $k_{B_C}$ | G-CSF binding rate | 2.24 | ml/pg/day |
| $k_{B_F}$ | IFN binding rate | 0.011 | ml/pg/day |
| $k_{U_L}$ | IL-6 unbinding rate | 22.3 | 1/day |
| $k_{U_G}$ | GM-CSF unbinding rate | 522 | 1/day |
| $k_{U_C}$ | G-CSF unbinding rate | 184 | 1/day |
| $k_{U_F}$ | IFN unbinding rate | 6.07 | 1/day |
| $POW$ | Stoichiometric constant (G-CSF)<br>Stoichiometric constant (IL-6, GM-CSF, IFN) | 1.4608<br>1 | Unitless |
| $\hat{p}$ | Stoichiometric constant (G-CSF)<br>Stoichiometric constant (IL-6, GM-CSF, IFN) | 2<br>1 | Unitless |

**Supplementary Table 9. Cytokine binding/unbinding rates and stoichiometric constant.**

| Parameter | Description | Value | Units |
|---|---|---|---|
| $K_{L,N}$ | No. IL-6 receptors on neutrophils | 720 | sites/cell |
| $K_{L,T}$ | No. IL-6 receptors on T cells | 300 | sites/cell |
| $K_{L,M}$ | No. of IL-6 receptors on monocytes | 509 | sites/cell |
| $K_{G,N}$ | No. of GM-CSF receptors on monocyte | 1058 | sites/cell |
| $K_{C,N}$ | No. of G-CSF receptors on neutrophil | 600 | sites/cell |
| $K_{F,T}$ | No. of IFN receptors on T cells | 1000 | sites/cell |
| $K_{F,I}$ | No. of IF receptors on infected cells. | 1300 | sites/cell |
| $MM_L$ | Molecular weight of IL-6 | 21000 | g/mol |
| $MM_G$ | Molecular weight of GM-CSF | 14000 | g/mol |
| $MM_C$ | Molecular weight of G-CSF | 19600 | g/mol |
| $MM_F$ | Molecular weight of IFN-$\beta$ | 19000 | g/mol |

**Supplementary Table 10. Number of cellular receptors and cytokine molecular weights.**

| Parameter | Description | Value | Units |
|---|---|---|---|
| $V_0$ | Initial viral load | 4.5 | log(copies/ml) |
| $S_0$ | Initial susceptible cells | 0.16 | $10^9$ cells/ml |
| $I_0$ | Initial infected cells | 0 | $10^9$ cells/ml |

| | | | |
|---|---|---|---|
| $R_0$ | Initial resistant cells | 0 | $10^9$ cells/ml |
| $M_{\Phi R,0}$ | Initial resident macrophages | $2.7\times 10^{-5}$ | $10^9$ cells/ml |
| $M_{\Phi I,0}$ | Initial inflammatory macrophages | $2.9\times 10^{-5}$ | $10^9$ cells/ml |
| $M_0$ | Initial monocytes | 0.0004 | $10^9$ cells/ml |
| $M_R$ | Initial reservoir monocytes | 0.0023 | $10^9$ cells/ml |
| $N_0$ | Initial neutrophils | 0.0053 | $10^9$ cells/ml |
| $N_R$ | Initial reservoir neutrophils | 0.0316 | $10^9$ cells/ml |
| $T_0$ | Initial CD8+ T cells | $1.1\times 10^{-4}$ | $10^9$ cells/ml |
| $L_{U,0}$ | Initial concentration of unbound IL-6 | 1.1 | pg/ml |
| $L_{B,0}$ | Initial concentration of bound IL-6 | $1.4\times 10^{-6}$ | pg/ml |
| $G_{U,0}$ | Initial concentration of unbound GM-CSF | 2.43 | pg/ml |
| $G_{B,0}$ | Initial concentration of bound GM-CSF | $1.6\times 10^{-6}$ | pg/ml |
| $C_{U,0}$ | Initial concentration of unbound G-CSF | 0.025 | pg/ml |
| $C_{B,0}$ | Initial concentration of bound G-CSF | $6.5\times 10^{-10}$ | pg/ml |
| $F_{U,0}$ | Initial concentration of unbound IFN | 0.015 | pg/ml |
| $F_{B,0}$ | Initial concentration of bound IFN | $1.1\times 10^{-8}$ | pg/ml |

**Supplementary Table 11. Initial conditions**

| Variable | Description | Units |
|---|---|---|
| $t$ | Time | days |
| $V$ | Viral load | cop/ml |
| $S$ | Number of susceptible cells | $10^9$ cells/ml |
| $I$ | Infected cells | $10^9$ cells/ml |
| $R$ | Resistant cells | $10^9$ cells/ml |
| $M_{\Phi R}$ | Alveolar (resident) macrophages | $10^9$ cells/ml |
| $M_{\Phi I}$ | Inflammatory macrophages | $10^9$ cells/ml |
| $M$ | Monocytes | $10^9$ cells/ml |
| $M_R$ | Bone marrow reservoir monocytes | $10^9$ cells/ml |
| $N$ | Neutrophils | $10^9$ cells/ml |
| $N_R$ | Bone marrow reservoir neutrophils | $10^9$ cells/ml |
| $T$ | CD8+ T cells | $10^9$ cells/ml |
| $L_U$ | Concentration of unbound IL-6 | pg/ml |
| $L_B$ | Concentration of bound IL-6 | pg/ml |
| $G_U$ | Concentration of unbound GM-CSF | pg/ml |
| $G_B$ | Concentration of bound GM-CSF | pg/ml |
| $C_U$ | Concentration of unbound G-CSF | pg/ml |
| $C_B$ | Concentration of bound G-CSF | pg/ml |
| $C_{BF}$ | Neutrophil G-CSF receptor bound fraction | unitless |
| $F_U$ | Concentration of unbound IFN | pg/ml |
| $F_B$ | Concentration of bound IFN | pg/ml |

**Supplementary Table 12. List of variables in the model equations.**

### Description of clinical data

Blood cell (monocytes, neutrophils, T cells) and cytokine (type-I IFN, IL-6, G-CSF, GM-CSF) measurements were collected from patients at the Centre hospitalier de l'Université de Montréal (CHUM) and the Jewish General Hospital in Montréal, Canada between March 2020 and May 2021. In this study, we used data from a total of 221 patients: 30 of whom were cancer patients and 24 of whom had immunosuppression (i.e., transplant recipients, patients receiving activated T-cell blockers like teriflunomide, or immunosuppressive drugs including Solu-Cortef and methylprednisolone).

To create the COVID-19 immunosuppressed virtual patient cohort, we used IL-6 concentrations measured on the 11$^{th}$ day after symptom onset from 19 immunosuppressed patients without autoimmune disease[2]. To generate the cancer virtual patient cohort, we calibrated the initial concentration of neutrophils to a study that included neutropenic patients with COVID-19[3], as cancer patients tend to have decreased neutrophil counts due to their anti-cancer treatments. Data were available from 23 COVID-19+ neutropenic patients who did not receive exogeneous G-CSF support, with measurements taken from 14 days prior to 14 days after symptoms onset; data collection times varied per study participants. To fit the cancer and reference virtual patients dynamics, CD8+ T cell data were incorporated from 93 COVID-19+ patients with cancer and 1959 COVID-19 patients without cancer[4]. T cells were measured on the 7$^{th}$ and 14$^{th}$ day after symptom onset.

### Comparisons of concentrations of IL-6, inflammatory macrophages, neutrophils, and damaged tissue to severity

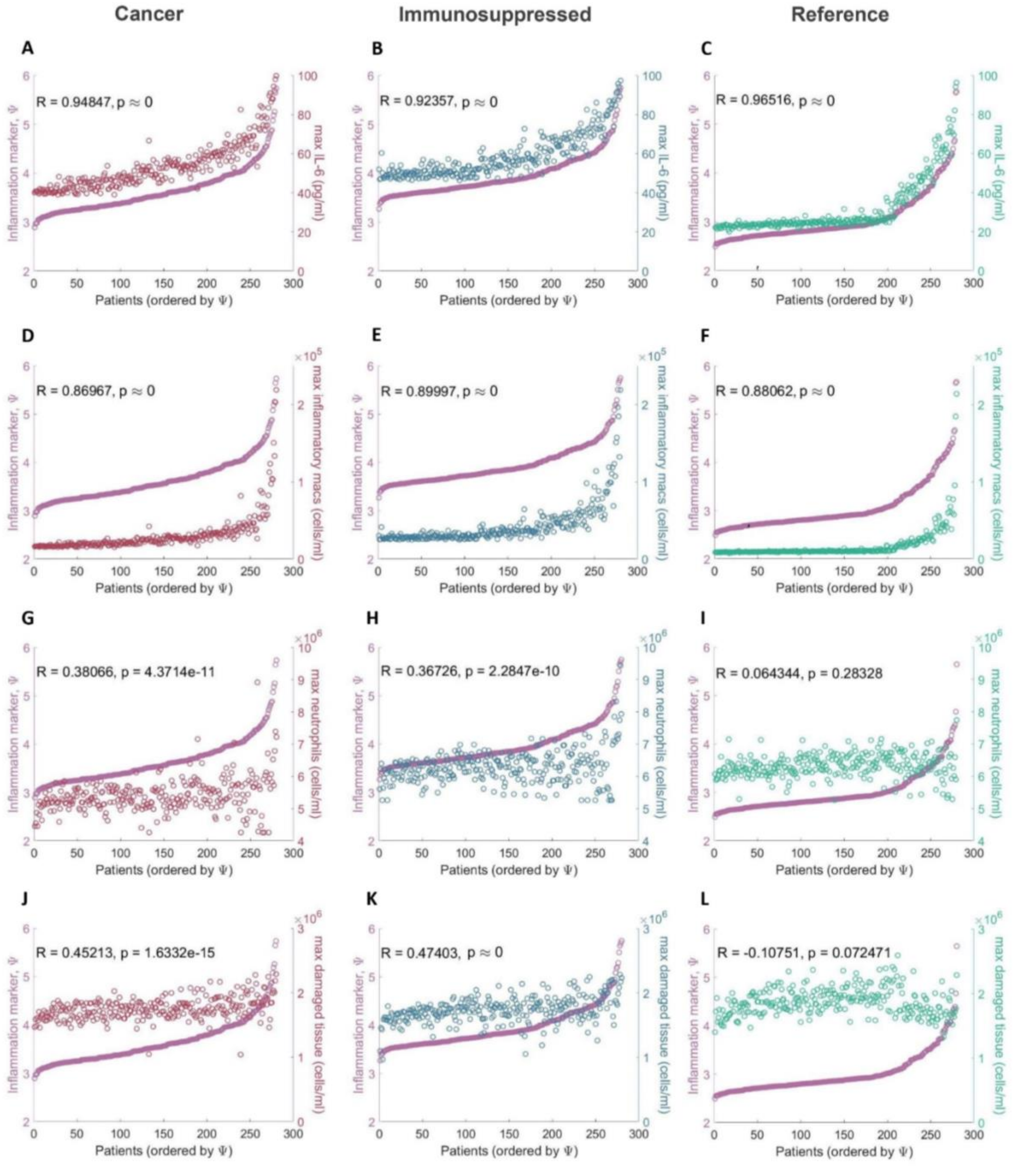


**Supplementary Figure 1. Comparisons of the maximal IL-6, inflammatory macrophage, and neutrophil concentrations, and maximal tissue damage with the inflammation marker.** Maximal IL-6 concentrations compared to the inflammation marker in patients from the A) COVID-19+ cancer cohort, B) COVID-19+ immunosuppressed cohort, and C) COVID-19+ reference cohort. Maximal IL-6 concentrations in all three cohorts were found to be positively correlated with $\Psi^j$. Maximal inflammatory macrophage values compared to the inflammation marker in patients from the D) COVID-19+ cancer cohort, E) COVID-19+ immunosuppressed cohort, and F) COVID-19+ reference cohort. Maximal inflammatory macrophages in all three cohorts were found to be positively correlated with $\Psi^j$. Maximal neutrophils compared to the inflammation marker in patients from the G)

COVID-19+ cancer cohort, H) COVID-19+ immunosuppressed cohort, and I) COVID-19+ reference cohort. There was no correlation between maximal neutrophils and the inflammation marker in either of the three cohorts. Maximal damaged tissue compared to the inflammation marker in patients from the J) COVID-19+ cancer cohort, K) COVID-19+ immunosuppressed cohort, and L) COVID-19+ reference cohort. There was no correlation between maximal damaged tissue and the inflammation marker in either of the three cohorts (p-values $< 10^{-10}$, indicating statistically significance).

**Statistical analyses of differences in parameter distributions between three virtual patient cohorts**

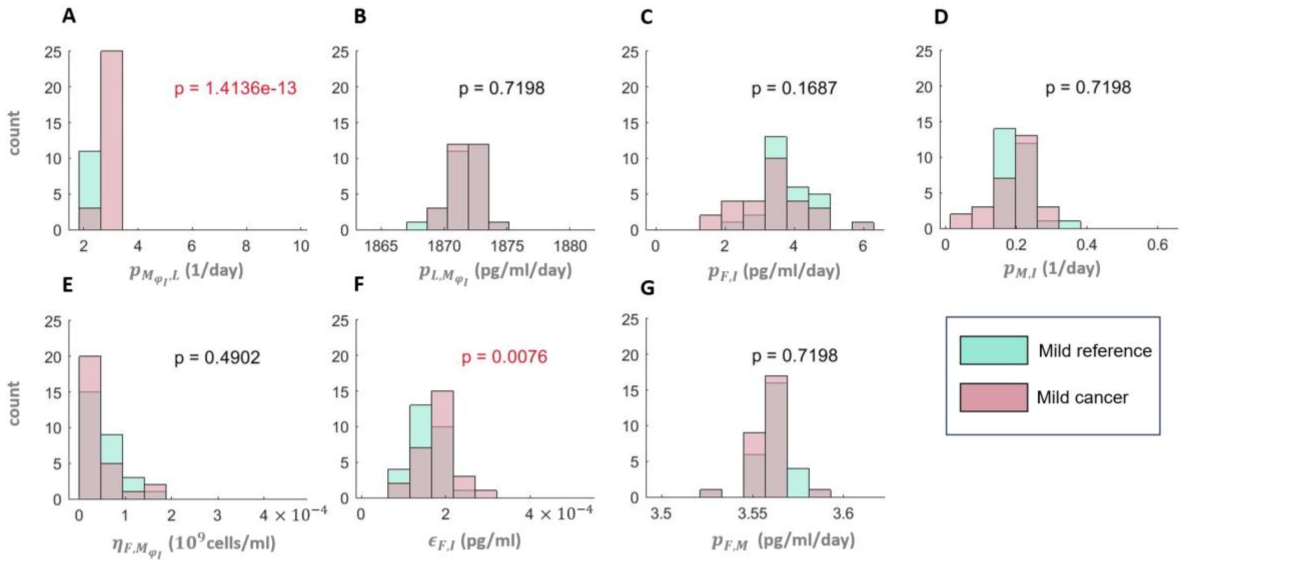


**Supplementary Figure 2. Parameter distribution comparisons between mild virtual patients in COVID-19+ cancer and COVID-19+ reference cohorts.** A) Monocyte-to-macrophage differentiation by IL-6, B) IL-6 production by inflammatory macrophages, C) IFN production rates by infected cells, D) Monocyte recruitment by infected cells, E) EC50 concentration of inflammatory macrophages on the IFN production, F) Cell-related IC50 concentration of IFN on virus production, and G) IFN production by monocytes. Statistically significant differences were found for parameters $p_{M_{\varphi_I},L}$ and $\epsilon_{F,I}$. Red p-values indicate statistically significant differences in distributions (p-value < 0.05).

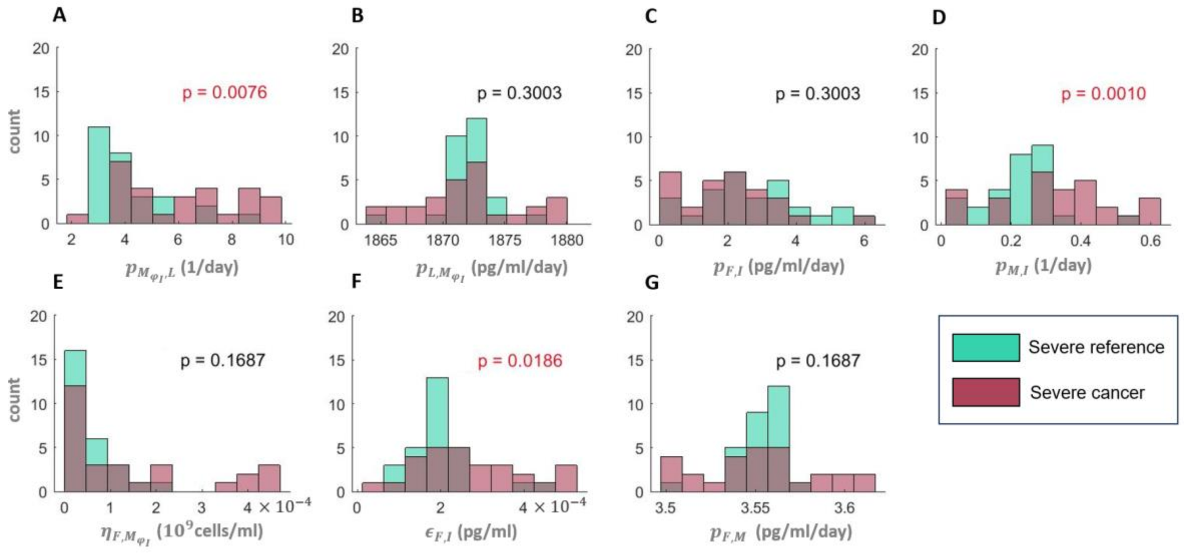


**Supplementary Figure 3. Parameter distribution comparisons between severe virtual patients in COVID-19+ cancer and COVID-19+ reference cohorts**. A) Monocyte-to-macrophage differentiation by IL-6, B) IL-6 production by inflammatory macrophages, C) IFN production rates by infected cells, D) Monocyte recruitment by infected cells, E) EC50 concentration of inflammatory macrophages on the IFN production, F) Cell-related IC50 concentration of IFN on virus production, and G) IFN production by monocytes. Statistically significant differences were found for parameters $p_{M_{\varphi_I},L}$, $p_{F,I}$, $p_{M,I}$, $\eta_{F,M_{\varphi I}}$, and $\epsilon_{F,I}$. Red p-values indicate statistically significant differences in distributions (p-value < 0.05).

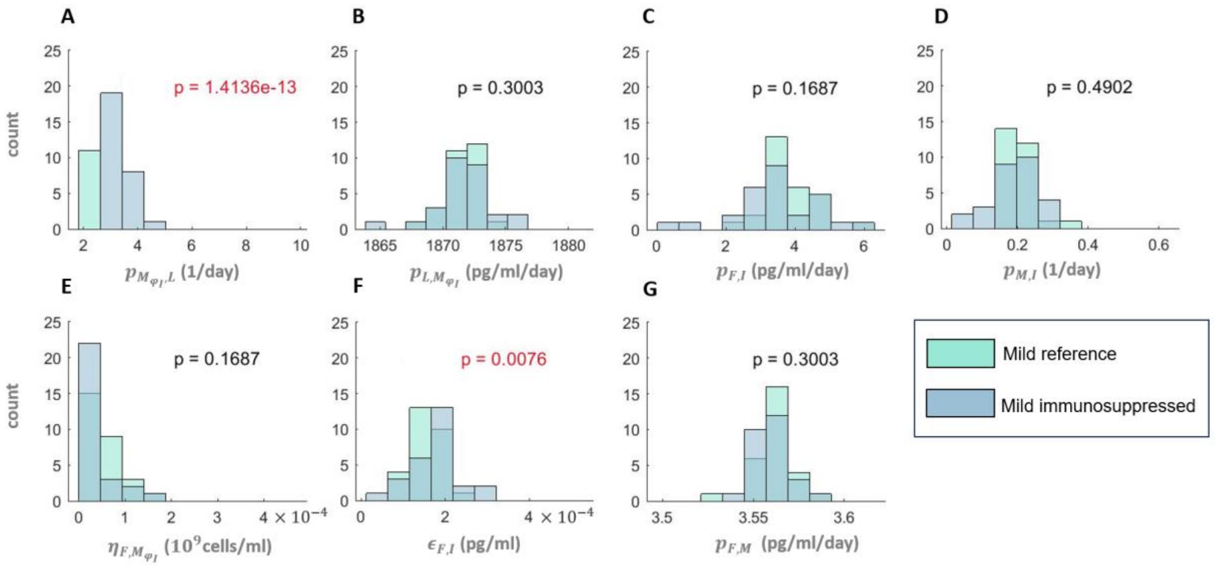


**Supplementary Figure 4. Parameter distribution comparisons between mild virtual patients in COVID-19+ immunosuppressed and COVID-19+ reference cohorts**. A) Monocyte-to-macrophage differentiation by IL-6, B) IL-6 production by inflammatory macrophages, C) IFN production rates by infected cells, D) Monocyte recruitment by infected cells, E) EC50 concentration of inflammatory macrophages on the IFN production, F) Cell-related IC50 concentration of IFN on virus production, and G) IFN production by monocytes. Statistically significant differences were found for parameters $p_{M_{\varphi_I},L}$ and $\epsilon_{F,I}$. Red p-values indicate statistically significant differences in distributions (p-value < 0.05).

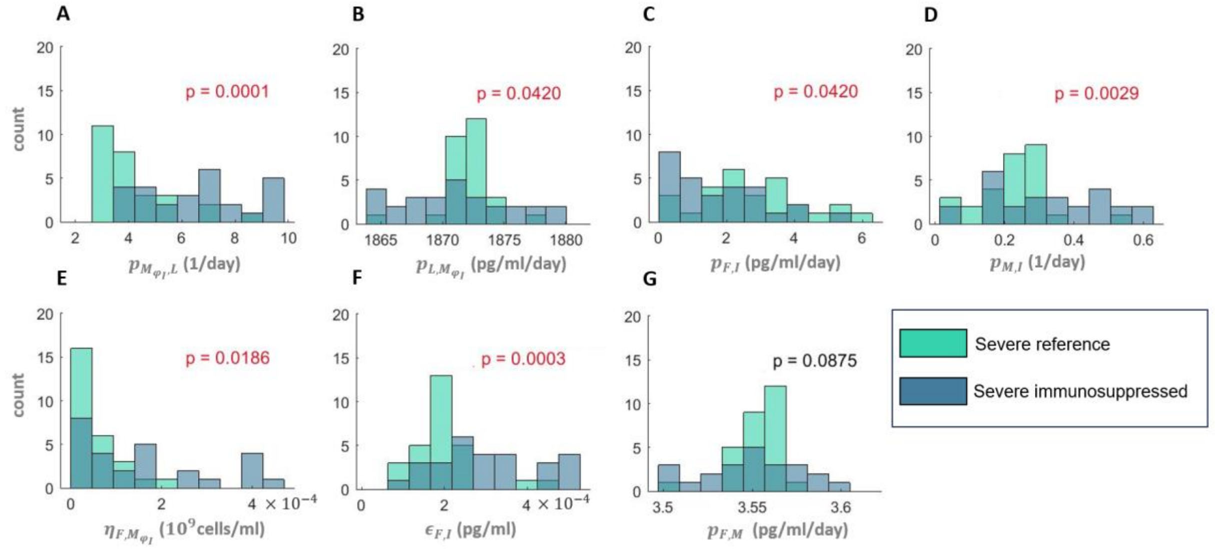


**Supplementary Figure 5. Parameter distribution comparisons between severe virtual patients in COVID-19+ immunosuppressed and COVID-19+ reference cohorts.** A) Monocyte-to-macrophage differentiation by IL-6, B) IL-6 production by inflammatory macrophages, C) IFN production rates by infected cells, D) Monocyte recruitment by infected cells, E) EC50 concentration of inflammatory macrophages on the IFN production, F) Cell-related IC50 concentration of IFN on virus production, and

G) IFN production by monocytes. Statistically significant differences were found for parameters $p_{M_{\varphi_I},L}$, $p_{L,M_{\varphi_I}}$, $p_{F,I}$, $p_{M,I}$, $\eta_{F,M_{\varphi I}}$, and $\epsilon_{F,I}$. Red p-values indicate statistically significant differences in distributions (p-value < 0.05).

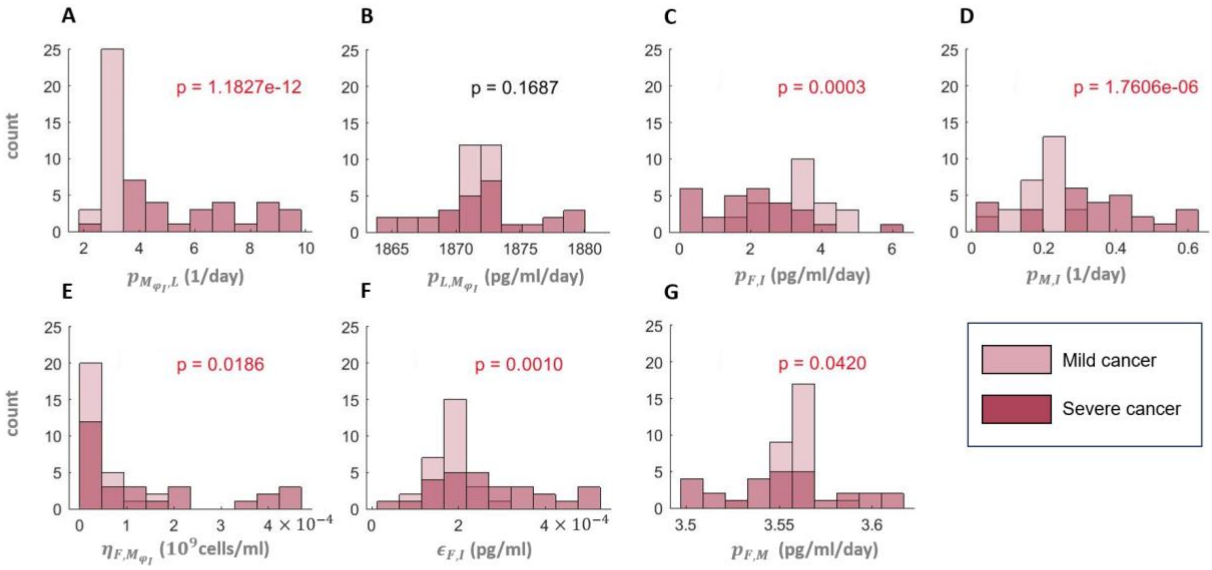


**Supplementary Figure 6. Parameter distribution comparisons between mild and severe virtual patients in the COVID-19+ cancer cohort**. A) Monocyte-to-macrophage differentiation by IL-6, B) IL-6 production by inflammatory macrophages, C) IFN production rates by infected cells, D) Monocyte recruitment by infected cells, E) EC50 concentration of inflammatory macrophages on the IFN production, F) Cell-related IC50 concentration of IFN on virus production, and G) IFN production by monocytes. Statistically significant differences were found for parameters $p_{M_{\varphi_I},L}$, $p_{F,I}$, $p_{M,I}$, $\eta_{F,M_{\varphi I}}$, $\epsilon_{F,I}$, and $p_{F,M}$. Red p-values indicate statistically significant differences in distributions (p-value < 0.05).

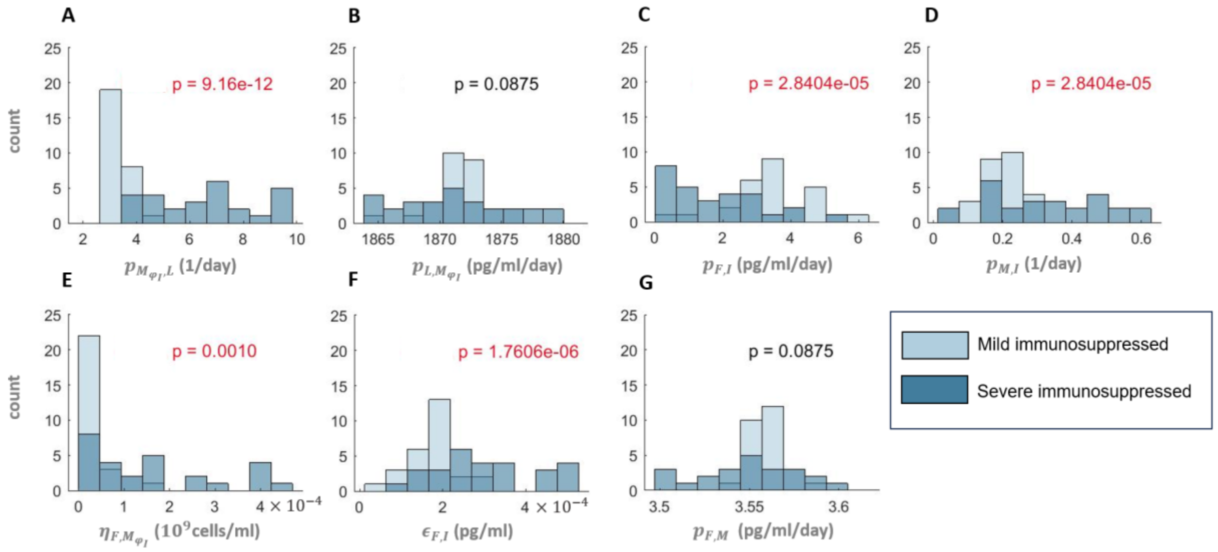


**Supplementary Figure 7. Parameter distribution comparisons between mild and severe virtual patients in the COVID-19+ immunosuppressed cohort.** A) Monocyte-to-macrophage differentiation by IL-6, B) IL-6 production by inflammatory macrophages, C) IFN production rates by infected cells, D) Monocyte recruitment by infected cells, E) EC50 concentration of inflammatory macrophages on the IFN production, F) Cell-related IC50 concentration of IFN on virus production, and G) IFN production by monocytes. Statistically significant differences were found for parameters $p_{M_{\varphi I},L}$, $p_{F,I}$, $p_{M,I}$, $\eta_{F,M_{\varphi I}}$, and $\epsilon_{F,I}$. Red p-values indicate statistically significant differences in distributions (p-value < 0.05).

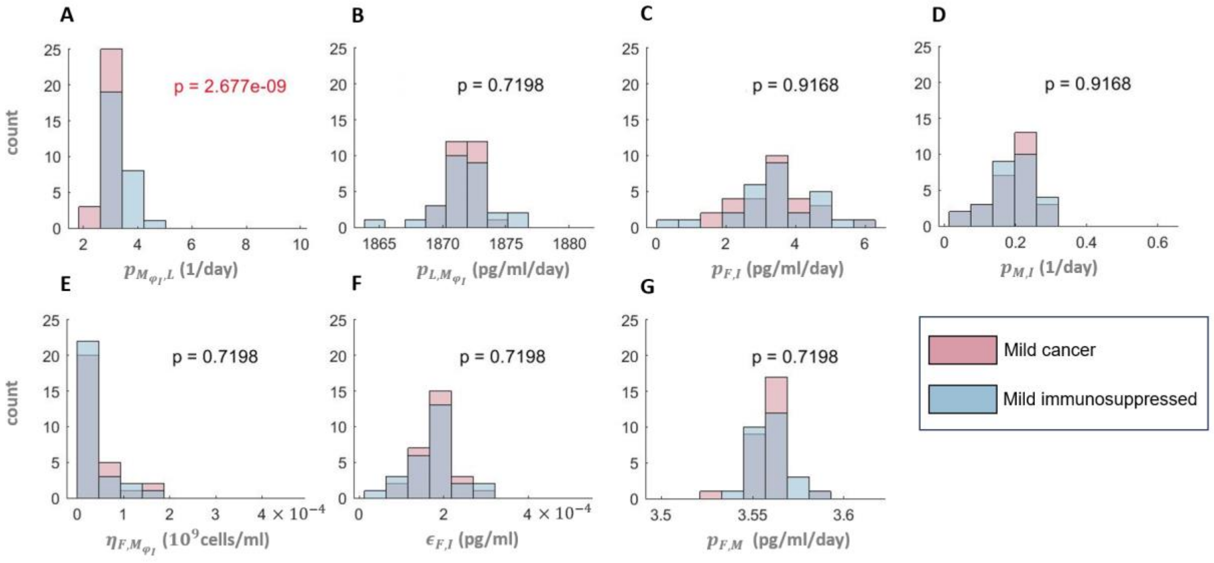


**Supplementary Figure 8. Parameter distribution comparisons between mild virtual patients in COVID-19+ cancer and COVID-19+ immunosuppressed cohorts**. A) Monocyte-to-macrophage differentiation by IL-6, B) IL-6 production by inflammatory macrophages, C) IFN production rates by infected cells, D) Monocyte recruitment by infected cells, E) EC50 concentration of inflammatory macrophages on the IFN production, F) Cell-related IC50 concentration of IFN on virus production, and G) IFN production by monocytes. Statistically significant differences were found for $p_{M_{\varphi_I},L}$. Red p-values indicate statistically significant differences in distributions (p-value < 0.05).

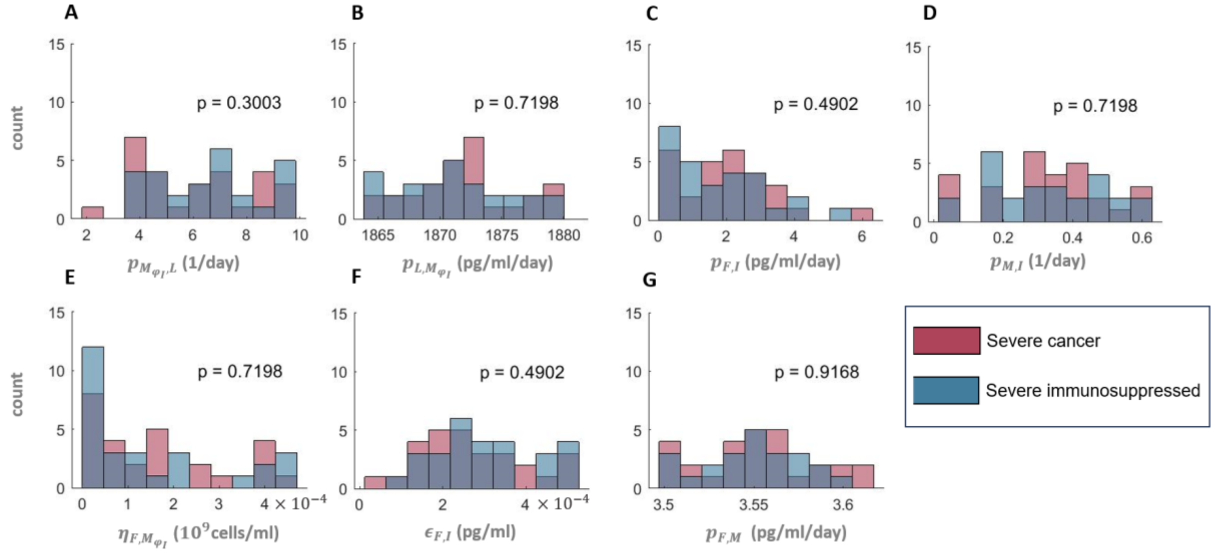


**Supplementary Figure 9. Statistical differences in parameter distributions between severe virtual patients in COVID-19+ cancer and COVID-19+ immunosuppressed cohorts.** A) Monocyte-to-macrophage differentiation by IL-6, B) IL-6 production by inflammatory macrophages, C) IFN production rates by infected cells, D) Monocyte recruitment by infected cells, E) EC50 concentration of inflammatory macrophages on the IFN production, F) Cell-related IC50 concentration of IFN on virus production, and G) IFN production by monocytes. Statistically significant differences were not found in any of the parameters. Red p-values indicate statistically significant differences in distributions (p-value < 0.05).

#### High IL-6 concentrations are not necessarily associated with delayed IFN peaks in cancer and immunosuppressed virtual patients

In Jenner et al.[1], we found a strong relationship between IFN and IL-6 where high maximal IL-6 concentrations were predictive of delayed IFN peak concentrations and thus severity. As expected, this connection was again established in our reference COVID-19 cohort. Interestingly, in this reference virtual patient cohort (VPC), we did not observe any delay in the time to IFN peak concentrations for virtual patients with maximum IL-6 concentrations below 55 pg/ml (Supplementary Figure 10Ac). However, in both the cancer and immunosuppressed VPCs, we identified several virtual patients with relatively low IL-6 concentrations who experienced delayed IFN peaks (Supplementary Figure 10Aa and 10Ab).

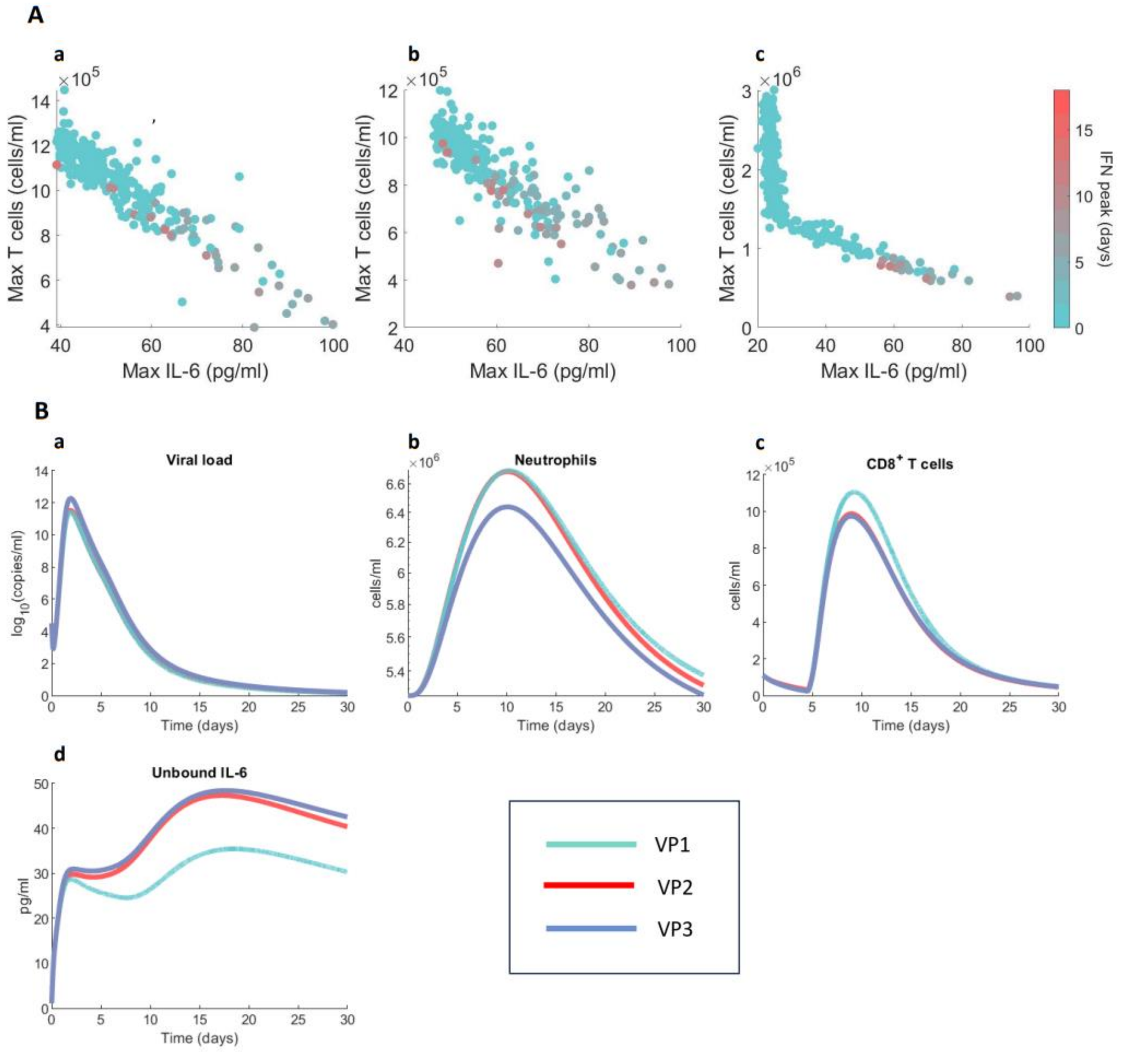


**Supplementary Figure 10. Relationships between maximum T cell and IL-6 concentrations, and IFN peak**. A: a) COVID-19+ cancer cohort, b) COVID-19+ immunosuppressed cohort, c) COVID-19+ reference cohort. B: a) Viral loads, b) Neutrophil concentrations, c) CD8+ T cells, and d) IL-6 concentrations in ‘extreme patients’ (VP1, VP2, and VP3).

**Features of outlier virtual patients reveal differences in parameters associated with IFN production**

To better understand the contrasting result of low IL-6 concentrations and IFN peaks, we studied the dynamics and parameter values of selected 'extreme' patients (VP1, VP2, and VP3) in the cancer VPC (Supplementary Figure 11A). Compared to other virtual patients in the cancer cohort, VP1 had the lowest peak IL-6 concentration, and VP2 and VP3 both had low maximum IL-6 and high maximal CD8+ T cell concentrations (Supplementary Figure 11A). These three virtual patients differed primarily in their IFN concentrations: VP2 had the highest peak IFN concentration while VP3 had the lowest IFN (Supplementary Figure 11B), neutrophils (Supplementary Figure 10Bb), and uninfected cell concentrations (Supplementary Figure 11C). VP1 had similar neutrophil and uninfected cell dynamics to VP2. Further, VP1 and VP2 had two pronounced IFN peaks, with the first arriving around the second day of infection (Supplementary Figure 11B), whereas the presence of two peaks was not observed in most other VPs.

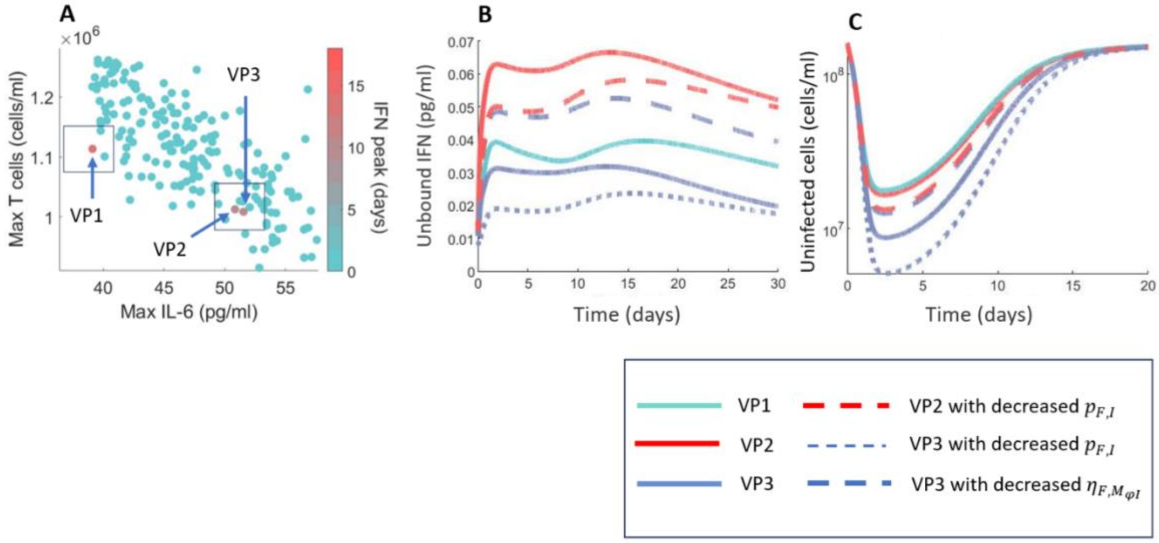


**Supplementary Figure 11. Parameter value changes in outlier virtual patients provide insight into mechanisms of immune dysregulation in cancer virtual patients.** A) Relationships between maximum IL-6, maximum T cell, and peak IFN concentrations in VP1, VP2, and VP3 in the COVID-19+ cancer cohort. B) Unbound IFN dynamics before and after decreasing values of parameters associated with IFN production. C) Uninfected cell dynamics of VP2 and VP3 before and after decreasing values of parameters associated with IFN production.

By analyzing VP1's, VP2's, and VP3's parameter values, we found that parameters related to IFN production (i.e., $\eta_{F,M_{\varphi I}}$ and $p_{F,I}$) most significantly differed between these three virtual patients (Supplementary Table 13). We next assessed the impact of these two parameter values on predicted immunological dynamics and COVID-19 severity by varying their values. VP3 had an $\eta_{F,M_{\varphi I}}$ that was about 3-fold higher than those of VP1 and VP2. Decreasing the value of $\eta_{F,M_{\varphi I}}$ in VP3 resulted in increased IFN concentrations and thus a higher number of uninfected cells (Supplementary Figure 11C). Similarly, VP2 had a $p_{F,I}$ value around 3 times larger than

VP1, and reducing its value caused a decrease in IFN and infected cell concentrations (Supplementary Figure 11B and 11C), see Discussion in the Main Text.

| | Parameter | | | | | | |
|---|---|---|---|---|---|---|---|
| | $p_{M_{\varphi I},L}$ | $p_{L,M_{\varphi I}}$ | $p_{F,I}$ | $p_{M,I}$ | $\eta_{F,M_{\varphi I}}$ | $\epsilon_{F,I}$ | $p_{F,M}$ |
| **VP1** | 2.47 | 1,874 | 0.13 | 0.26 | $1.82\times10^{-5}$ | $1.13\times10^{-4}$ | **3.57** |
| **VP2** | 2.88 | 1,872 | 0.35 | 0.26 | $1.08\times10^{-5}$ | $2.08\times10^{-4}$ | **3.56** |
| **VP3** | 3.05 | 1,872 | 0.33 | 0.22 | $7.30\times10^{-5}$ | $2.17\times10^{-4}$ | **3.58** |

**Supplementary Table 13. Parameter values of 'extreme' patients.** Shaded values of parameters that differed the most were decreased by 3 times ($p_{F,I}$ in VP3 and VP2) and 7 times ($\eta_{F,M_{\varphi I}}$ in VP3) their original values to see the impact of that change on biomarkers dynamics and thus severity.

### Predicted damaged tissue and IFN dynamics

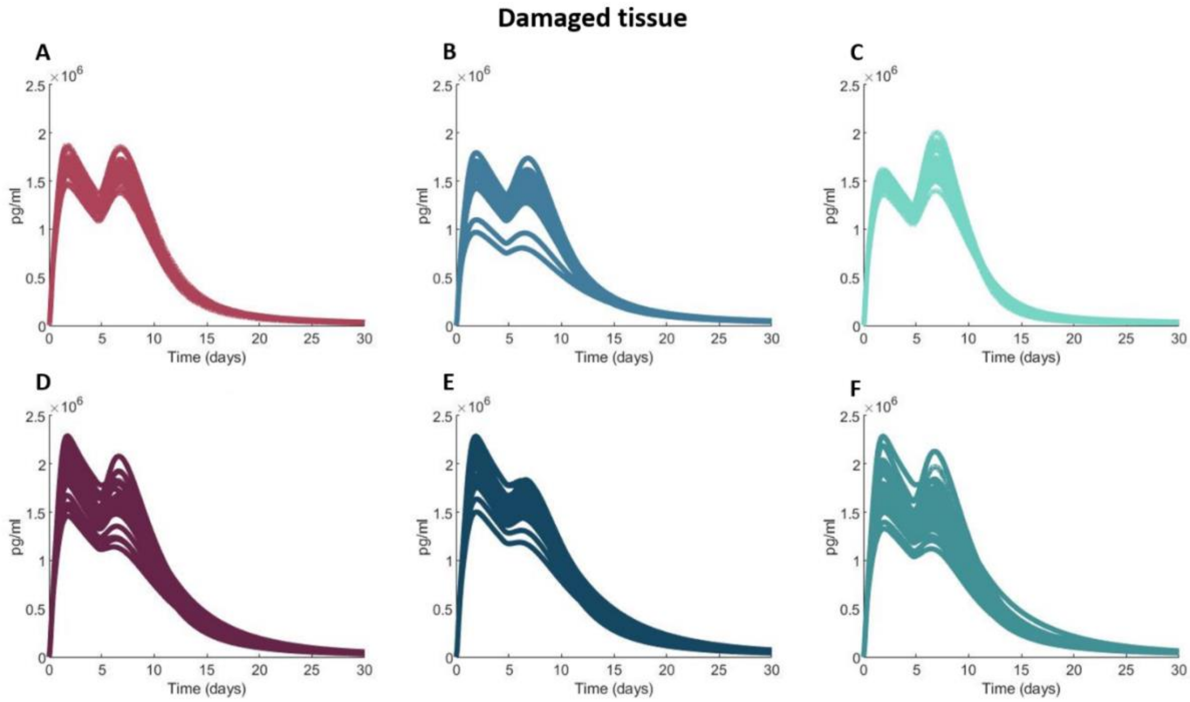


**Supplementary Figure 12. High tissue damage in severe patients in all three cohorts.** Dynamics of damaged tissue in A) mild COVID-19+ cancer patients, B) mild COVID-19+ immunosuppressed patients, C) mild COVID-19+ patients, D) severe COVID-19+ cancer patients, E) severe COVID-19+ immunosuppressed patients, F) severe COVID-19+ patients.

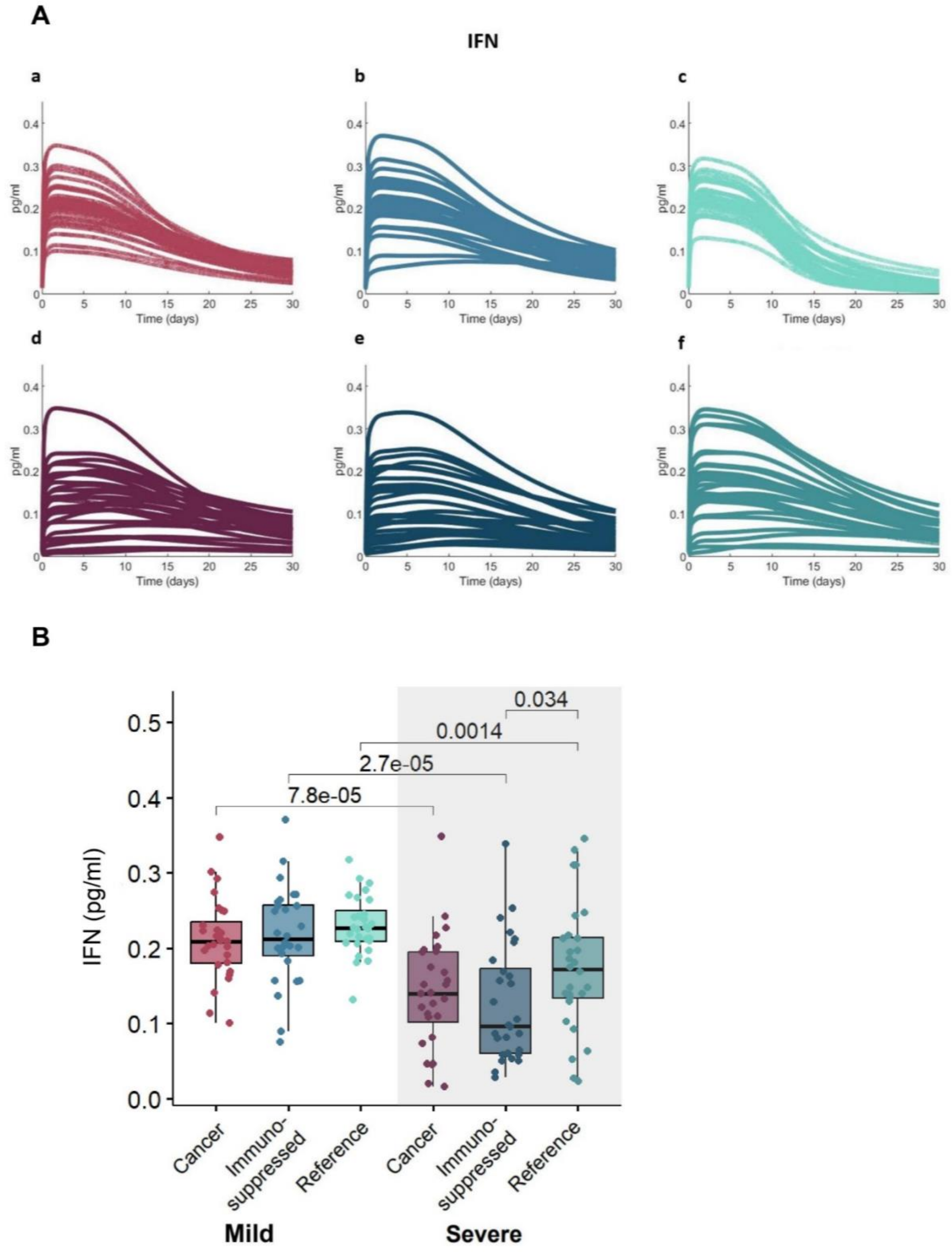


**Supplementary Figure 13. Maximum IFN concentrations are decreased in severe patients.** Dynamics of IFN in A: a) Mild COVID-19+ cancer patients, b) Mild COVID-19+ immunosuppressed patients, c) mild COVID-19+ patients, d) Severe COVID-19+ cancer patients, e) Severe COVID-19+ immunosuppressed patients, f) Severe COVID-19+ patients. B) Model predictions of mean values of maximal IFN concentrations in mild and severe virtual patients. Statistical differences in maximal IFN values between groups are indicated by the p-values above the boxplots for p-values < 0.05.
